# Metabolic fingerprinting reveals roles of *Arabidopsis thaliana* BGLU1, BGLU3 and BGLU4 in glycosylation of various flavonoids

**DOI:** 10.1101/2024.01.30.577901

**Authors:** Jana-Freja Frommann, Boas Pucker, Lennart Malte Sielmann, Caroline Müller, Bernd Weisshaar, Ralf Stracke, Rabea Schweiger

## Abstract

Flavonoids are specialized metabolites that play important roles in plants, including interactions with the environment. The high structural diversity of this metabolite group is largely due to enzyme-mediated modifications of flavonoid core skeletons. In particular, glycosylation with different sugars is very common. In this study, the functions of the *Arabidopsis thaliana* glycoside hydrolase family 1-type glycosyltransferase proteins BGLU1, BGLU3 and BGLU4 were investigated, using a reverse genetics approach and untargeted metabolic fingerprinting. We screened for metabolic differences between *A. thaliana* wild type, loss-of-function mutants and overexpression lines and partially identified differentially accumulating metabolites, which are putative products and/or substrates of the BGLU enzymes. Our study revealed that the investigated BGLU proteins are glycosyltransferases involved in the glycosylation of already glycosylated flavonoids using different substrates. While BGLU1 appears to be involved in the rhamnosylation of a kaempferol diglycoside in leaves, BGLU3 and BGLU4 are likely involved in the glycosylation of quercetin glycosides in *A. thaliana* seeds. In addition, we present evidence that BGLU3 is a multifunctional enzyme that catalyzes other metabolic reactions with more complex substrates. This study deepens our understanding of the metabolic pathways and enzymes that contribute to the high structural diversity of flavonoids.

**Highlight:** The proteins BGLU1, BGLU3 and BGLU4 are involved in glycosylations of different, already glycosylated flavonoids in *Arabidopsis thaliana*. BGLU3 appears to be multifunctional, acting on several complex substrates.

## Introduction

Plant specialized metabolites are of great importance for plant functioning (Kessler and Kalske, 2018). One of the best studied groups of these metabolites are flavonoids (Tohge *et al*., 2013; Wen *et al*., 2020). Flavonoids are characterized by a C6-C3-C6 core skeleton, with two aromatic rings (A and B) that are linked with a three-carbon bridge (C ring). They include various subclasses such as flavonols, anthocyanidins and flavanols which have distinct substitution patterns. Among the multiple functions that flavonoids fulfill in plants is the protection against diverse abiotic and biotic stresses, especially due to their antioxidative activities. They also attract beneficial organisms such as pollinators and seed dispersers via the generation or modification of flower and fruit colors (Falcone Ferreyra *et al*., 2012; Le Roy *et al*., 2016; Mierziak *et al*., 2014). The plethora of flavonoid functions is thought to be linked to their high structural diversity, with more than 8,000 known compounds and probably many more that have not yet been described (Tohge *et al*., 2013; Wen *et al*., 2020). While the core flavonoid biosynthetic pathway is well understood, knowledge about flavonoid-modifying enzymes is limited and incomplete (Tohge *et al*., 2013; Wen *et al*., 2020).

Specific enzymes modify the flavonoid skeleton (Le Roy *et al*., 2016; Tanaka *et al*., 2008), with glycosylation being one of the most abundant modifications. Sugar moieties, commonly glucose, galactose, rhamnose, xylose, arabinose and/or glucuronic acid, are bound to the skeleton by *O*-or *C*-glycosidic linkages (Kachlicki *et al*., 2016; Noguchi *et al*., 2009). Because sugar residues enhance the water solubility, protect reactive nucleophilic groups and increase metabolite stability (Gachon *et al*., 2005; Plaza *et al*., 2014), glycosylation is important for metabolite transport and storage (Le Roy *et al*., 2016; Wang *et al*., 2019). Oligo-/polymerization and condensation reactions further increase the structural diversity of flavonoids. These condensates include oligo-/polymers of flavanols like proanthocyanidins or condensed tannins (Dixon *et al*., 2005), condensates of anthocyanins with flavanols (Gonzalez-Manzano *et al*., 2008; Lee *et al*., 2009; Remy *et al*., 2000) and pyranoanthocyanins that are formed by reactions of anthocyanins with small non-flavonoid molecules and/or other flavonoids to form a new pyrane ring (Andersen *et al*., 2004; Fulcrand *et al*., 1998; Nave *et al*., 2010; Rentzsch *et al*., 2007).

Glycosylation is catalyzed by glycosyltransferases (GTs), most of them utilizing an uridine diphosphate (UDP)-activated sugar as donor, which leads to the designation UDP-GTs or UGTs (Le Roy *et al*., 2016). Moreover, acyl-glucose-dependent GTs (AGTs) glycosylate already substituted flavonoids at the skeleton or substituents. Such sugar transfers to anthocyanin acceptors have been described for different plant species, including *Arabidopsis thaliana* (Brassicaceae) (Matsuba *et al*., 2010; Miyahara *et al*., 2013; Miyahara *et al*., 2012). Evidence for transfer of acyl-derived sugars to flavonol glucoside acceptors has, to our knowledge, so far only been provided for *A. thaliana* (Ishihara *et al*., 2016). The AGTs exhibit high amino acid sequence similarity to glycoside hydrolase family 1 (GH1) proteins that typically act as *β*-glucosidases (BGLUs) (Matsuba *et al*., 2010; Miyahara *et al*., 2012). Phylogenetic analysis of 47 BGLUs of *A. thaliana* revealed a cluster of BGLU1 to BGLU11 (Miyahara *et al*., 2011; Xu *et al*., 2004), including the potential AGTs BGLU6 (Ishihara *et al*., 2016) and BGLU10 (Miyahara *et al*., 2013). Since phylogenetic clustering might suggest similar protein functions, BGLU1 to BGLU11 may all act as AGTs (Miyahara *et al*., 2011), while the separation of BGLU1 to BGLU6 within this cluster points to an activity on non-anthocyanin flavonoid substrates, as genetically demonstrated for BGLU6 (Ishihara *et al*., 2016).

In the present study, we investigated the functions of the *A. thaliana BGLU* genes *BGLU1*, *BGLU3* and *BGLU4*, which cluster around *BGLU6*. We used a reverse genetics approach based on *bglu* loss-of-function mutants and *BGLU* overexpression lines. Metabolic differences between these expression variant lines were investigated by untargeted metabolic fingerprinting. We present evidence that the enzymes encoded by *BGLU1*, *BGLU3* and *BGLU4* show GH1-type GT, probably AGT, activity in *A. thaliana* leaves (*BGLU1*) and seeds (*BGLU3*, *BGLU4*), acting on glycosylated flavonols as substrates. BGLU3 appears to be a multifunctional GT, since several putative products and substrates of this enzyme were detected.

## Materials and methods

### Plant material

*A. thaliana* seeds were obtained from the Nottingham Arabidopsis Stock Center, including the wild type (wt) Columbia-0 (Col-0) accession and the loss-of-function transfer DNA (T-DNA) insertion mutants *bglu1-1* (*At1g45191*, GK_341B12, N432664), *bglu3-2* (*At4g22100*, GK_853H01, N481877; Kleinboelting *et al*., 2012) and *bglu4-2* (*At1g60090*, SALK_029729, N25045; Alonso *et al*., 2003). Homozygous plants were selected based on PCR-based genotyping (Supplementary Table S1 at *JXB* online) and kept as mutant lines by selfing. The insertion alleles and genomic integrity of the T-DNA insertion mutants were verified by long-read sequencing (see below). Overexpression lines, based on the double enhancer cauliflower mosaic virus 35S (2x35S) promoter (Kay *et al*., 1987), were generated in Col-0 wt as described below.

### Plant growth conditions

Unless stated otherwise, plants were grown in the greenhouse under long day conditions (about 14 h light) at 23 °C and 70% relative humidity (rH). They were grown in 9 x 9 cm pots on compost “Sondermischung Max Planck Institut 19277634” consisting of 70% white peat (finely ground), 20% vermiculite^®^ and 10% sand with pH 6.5; the substrate contained 1 kg m^-^³ Osmocote^®^ Start 8 Weeks (starting fertilizer); 1 kg m^-^³ Triabon^®^ (depot fertilizer) and 0.25 kg m^-^³ Fe-EDDHA. Plants were fertilized with Wuxal^®^ Super (Manna) and watered as required. Until flowering, pots of the different genotypes were placed in random order. To prevent cross-pollination at the flowering stage, pots of the same genotype were placed in trays that were randomized daily. For seed formation, plants were grown under short day conditions (8 h light, 22 °C, about 55% rH) for two months before being transferred to long day conditions. For seed ripening, the temperature was raised to 24 °C. Leaf samples for gene expression and metabolic studies were obtained from 6-week-old plants, which were grown for 8 days on 0.5x Murashige and Skoog medium, followed by 5 weeks of growth in a growth chamber (Percival) with 13 h light at 24 °C and 80% rH (18 °C and 65% rH at night).

### Long-read sequencing of genomic DNA

For long-read sequencing of homozygous *bglu1-1*, *bglu3-2* and *bglu4-2* mutants, high molecular weight genomic DNA was extracted from young rosette leaves and analyzed as described (Pucker *et al*., 2021). Library preparation followed the SQK-RAD004 (*bglu3-2*) or SQK-LSK109 (*bglu1-1, bglu4-2*) protocol (Oxford Nanopore Technologies) and sequencing was performed on R9.4.1 and R10 flow cells (Supplementary Table S2). The sequencing data were submitted to the *European Nucleotide Archive under study ID PRJEB36305*: *bglu1-1* (ERS4255859), *bglu3-2* (ERS4255860) and *bglu4-2* (ERS4255861). The identified T-DNA::genome junctions were confirmed by Sanger sequencing of PCR amplicons.

### RNA isolation and cDNA synthesis

Informed by RNA-Seq-derived expression data from the TraVa website (Klepikova *et al*., 2016), RNA was isolated from plant parts with the highest *BGLU* expression in Col-0 wt. Around 10 mm long rosette leaves were used for *BGLU1* (813 read counts by median normalization, see TraVa), dry mature seeds for *BGLU3* (4,006 read counts) and mature seeds soaked in water on a wet filter paper for 24 h in the dark for *BGLU4* (4,062 read counts). Seed samples were taken from multiple plants grown in the same batch. RNA isolation was performed using the Spectrum™ Plant Total RNA Kit (Protocol A; Sigma-Aldrich) according to the supplier‘s instructions, including DNase I digestion. Seed RNA samples for RT-PCR were precipitated with 8 M LiCl as described (Suzuki *et al*., 2004). cDNA was synthesized from 1 µg total RNA using the ProtoScript^®^ II First Strand cDNA Synthesis Kit (NEB) with d(T)_23_VN primer.

### cDNA cloning

Full-length coding sequence (CDS) constructs of *BGLU1*, *BGLU3* and *BGLU4* were created using the GATEWAY^®^ Technology (Invitrogen). PCRs were performed with the Q5^®^ High Fidelity DNA Polymerase (NEB) and specific, *attB* recombination site-containing primers (Supplementary Table S1). The resulting amplicons were recombined into the GATEWAY^®^ vector pDONR™/Zeo with BP clonase resulting in entry constructs. All constructs were verified by Sanger sequencing.

### Generation of BGLU overexpression lines

The full-length *BGLU1*, *BGLU3* or *BGLU4* CDSs from the entry constructs were introduced into the binary expression vector pLEELA (Jakoby *et al*., 2004) using GATEWAY^®^ LR clonase. The T-DNA from the resulting plasmid constructs with *2x35S::BGLU::35S-polyA* expression cassettes was transferred into *A. thaliana* via *Agrobacterium tumefaciens*-mediated [Agrobacterium, GV101::pMP90RK; Koncz and Schell, 1986] gene transfer by floral dip (Clough and Bent, 1998). Positive lines were identified by BASTA-selection and confirmed by PCR-based genotyping. Transgene expression was analyzed in rosette leaves for the *2x35S::BGLU1*, *2x35S::BGLU3* and *2x35S::BGLU4* lines by RT-PCR to select lines with the highest *BGLU* transcript levels.

### Differential gene expression of BGLU1, BGLU3 and BGLU4

Gene expression for *BGLU1*, *BGLU3* and *BGLU4*, respectively, was examined in biological triplicates (*BGLU1*: leaves of single plants, *BGLU3*/*BGLU4*: seeds from several plants) by reverse transcriptase quantitative real-time PCR (RT-qPCR). Sample collection for RNA isolation was performed as described for the metabolic analyses (see below). Intact transcripts of the *bglu* mutants were quantified using intron-spanning amplimers, with primers designed using the Python script *find_primers.py* (https://github.com/hschilbert/Primer_design). RT-qPCR was performed in technical triplicates using the Luna^®^ Universal qPCR Master Mix (NEB) in a CFX96 Touch™ Real-Time PCR Detection System (Bio-Rad, 39 cycles).

Transcript levels were quantified relative to the wt, using the 2^−ΔΔCt^ method (Livak and Schmittgen, 2001) and the geometric mean of the reference genes *PEROXIN4* (*At5g25760*) and *EF1α* (*At5g60390*) (Vandesompele *et al*., 2002). The additive error was calculated for the asymmetrically distributed standard errors relative to the mean. If C_t_ values in the *bglu* samples were zero, the maximum cycle number plus one (Ct = 40) was taken for calculations. To compare the gene expression levels with the intensities of metabolic features that may represent products and/or substrates of the enzymes encoded by the investigated genes (metabolic fingerprinting, see below), reverse transcriptase semi-quantitative PCR (RT-PCR) was applied as fast approach. This was done with the plants that were used for metabolic analyses and with *ACTIN2* (*At3g18780*) as reference gene.

### Subcellular localization of BGLU-RFP fusion proteins

Full-length *BGLU1*, *BGLU3* or *BGLU4* CDSs were introduced into the binary expression vector pUBC-RFP-Dest (Grefen *et al*., 2010) using GATEWAY^®^ LR clonase, resulting in *pUBQ10::BGLU::RFP* fusion constructs. *2x35S::GFP* in pAVA393 (Nesi *et al*., 2001) was used as control for cytoplasmic and nuclear localization (von Arnim *et al*., 1998). TRANSPARENT TESTA13 (TT13)-GFP (*p35S::TT13-GFP* in pK7FWG2) (Appelhagen *et al*., 2015) was used as an endomembrane control for vacuolar localzation. The constructs were (co-)transfected into tobacco Bright Yellow-2 (BY-2) protoplasts as previously described (Haasen *et al*., 1999), using 30 µg (single transfection) or 20 µg (co-transfection) DNA of each plasmid. RFP and GFP fluorescence were detected after 24 h of incubation in the dark using an inverted confocal laser scanning microscope 780 (LSM 780, Zeiss) with a water-immersion oil objective (LCI Plan-Neofluar 63x /1.3 Imm Korr DIC M27, Zeiss) and the main beam splitter MBS488/561. An argon ion laser at 488 nm (GFP) or a diode-pumped solid-state laser at 561 nm (RFP) was used for excitation and detection at 493–551 nm (GFP) or 582–702 nm (RFP), respectively. Images were acquired with a pixel dwell time of ≤ 6.3 µs, an intensity resolution of 12 or 16 bit per pixel, an 8-to 16-fold averaging (depending on noise) and adjusted similar maximal brightness in both channels to correct photostability and brightness differences of GFP and RFP. Images were processed using ZEN (2011, Zeiss) and Fiji (ImageJ) v2.0.0 (Schindelin *et al*., 2012). Co-localization was visualized with merged images, where pixels with positive signals for RFP and GFP are shown in white as described (Dunn *et al*., 2011).

### Untargeted metabolic fingerprinting

Metabolic fingerprinting was used to screen for candidate product and substrate metabolites of the biosynthetic reactions catalyzed by the investigated enzymes. For each investigated *BGLU* gene, a sample set containing (i) the *bglu* loss-of-function mutant (*bglu1-1*, *bglu3-2* or *bglu4-2*), (ii) the wt and (iii) the overexpression line (*2x35S::BGLU1* #6, *2x35S::BGLU3* #44 or *2x35S::BGLU4* #91) was prepared. The plant part with the highest *BGLU* gene expression in the wt (see above) was used for the metabolic analyses. For BGLU1 samples, four to six rosette leaves of 6-week-old plants were harvested 6 h after artificial sunrise. For *BGLU3* samples, 150 mg dry mature seeds were used, and for *BGLU4* samples 120 mg seeds, which were sown on wet filter paper to soak with water for 24 h in the dark (all seeds were 5 to 6 months old, derived from a pool of plants). Four biological replicates were prepared for each genotype. Samples were flash frozen in liquid nitrogen, stored at −80 °C, lyophilized and ground. Extraction and analysis of (semi)-polar metabolites were performed as described (Schrieber *et al*., 2019) with some modifications. Samples (10 mg powder) were extracted threefold in ice-cold 90% (v:v) methanol (LC-MS grade; Fisher Scientific UK Limited or Th. Geyer GmbH & Co. KG), supplemented with luteolin 7-*O*-glucoside (Extrasynthese) as internal standard. Pooled supernatants were filtered using Phenex™ syringe filters (0.2 µm, Phenomenex^®^). One blank was prepared for each set of ten samples.

Samples were analyzed using an ultra-high performance liquid chromatograph (Dionex UltiMate 3000, Thermo Fisher Scientific) and a quadrupole time-of-flight mass spectrometer (compact, Bruker Daltonics) in positive electrospray ionization (ESI^+^) mode. Separation was done on a Kinetex XB-C18 column (1.7 µm, 150 mm x 2.1 mm, with guard column; Phenomenex) at 45 °C with a flow rate of 0.5 ml min^-1^. As mobile phases, 0.1% (v:v) formic acid (∼98%, LC-MS grade, Honeywell Research Chemicals, Fluka) in H_2_O_MilliQ_ (phase A) and 0.1% formic acid in acetonitrile (LC-MS grade; Fisher Scientific or HiPerSolv CHROMANORM, VWR) (phase B) were used, with a gradient increasing linearly from 2% to 30% B within 20 min and to 75% B within 9 min, followed by column cleanup and equilibration. A nebulizer (N_2_) pressure of 3 bar, an end plate offset of 500 V, a capillary voltage of 4,500 V and N_2_ as drying gas (275 °C, flow rate: 12 l min^-1^) were used. A Na(HCOO)-based calibration solution was introduced to the ESI sprayer before or after each sample. Line mass spectra were recorded in the mass-to-charge (*m*/*z*) range of 50–1,300 *m*/*z* at 1-8 Hz, depending on the type of sample (plant part) and peak heights; the same spectra rate was used for samples to be compared (see below). The MS parameters were: 4 eV quadrupole ion energy, a low mass with an *m*/*z* value of 90, 7 eV collision energy, 75 µs transfer time and 6 µs pre-pulse storage. To obtain fragment (MS/MS) spectra of the ions with the highest intensities, the Auto-MS/MS mode was used with N_2_ as collision gas and the isolation widths and collision energies increased with the *m*/*z* of the precursors. To aid in metabolite identification, some samples were additionally measured at low spectra rates (1–3 Hz) and using multiple reaction monitoring to specifically fragment certain ions, sometimes using different collision energies. Some samples were also measured in negative electrospray ionization mode (ESI^−^; capillary voltage 3,000 V) for aglycone identification; since these measurements were performed at different time points, the retention times in ESI^+^ and ESI^−^ modes slightly differ.

Mass axis recalibration using the Na(HCOO) calibrant and picking of metabolic features [each characterized by a retention time (RT) and *m*/*z*] including spectral background subtraction were performed in Compass DataAnalysis v4.4 (Bruker Daltonics). The “Find Molecular Features” algorithm of the Bruker DataAnalysis software was used for feature picking, with the following settings: signal-to-noise threshold 3 (or 1 if measured at 1 Hz), correlation coefficient threshold 0.75; depending on the spectra rates, minimum compound lengths were set to 5-22 spectra and smoothing widths to 0-6. Using Compass ProfileAnalysis v2.3 (Bruker Daltonics), metabolic features likely to belong to the same metabolite (i.e., [M+H]^+^ ions, common adducts and fragments with corresponding isotopes and charge states) were grouped together in so-called buckets. The split-buckets-with-multiple-compounds option was selected to separate the internal standard from a peak with similar RT and *m*/*z* in seed samples. Each bucket was reduced to the feature with the highest intensity in that bucket and this feature was used for quantification via its peak height. These features were aligned across samples, allowing RT deviations of 0.1 or 0.2 min and *m*/*z* deviations of 6 mDa, respectively. Features within the injection peak and those with peak heights above detector saturation were excluded. Peak heights were related to the height of the [M+H]^+^ ion of the internal standard. Based on the resulting values, features were retained in the data set of the corresponding gene, if their mean intensity in at least one genotype of a sample set (*bglu* mutant, wt, *2x35S::BGLU* line) was at least 50 times higher than the corresponding intensity in the blanks. Moreover, features had to be present in at least three of the four biological replicates in at least one genotype. Finally, the feature intensities were divided by the sample dry weight.

### Screening for and (partial) identification of metabolites

To screen for metabolites that may represent products and substrates of the enzymes encoded by *BGLU1*, *BGLU3* and *BGLU4*, fold changes (FCs) were calculated. For this, the mean intensities of all metabolic features in the *BGLU* expression variant lines were divided by the corresponding mean intensities in the wt if at least one genotype in the pairwise combination showed peaks in at least three replicates. Candidate product features were selected based on higher peaks in *2x35S::BGLU* than in wt samples (FC ≥ 1.5 or present only in *2x35S::BGLU* but not in wt samples) and/or lower peaks in *bglu* than in wt samples (FC ≤ 0.67 or only occurring in wt but not in *bglu* samples). Candidate substrate features were selected by screening for the opposite peak intensity patterns. Extracted ion chromatograms of the *m*/*z* belonging to the features of interest were manually reviewed and features were considered relevant if the peak intensity patterns across genotypes resembled the transcript expression patterns from the RT-PCR (candidate products) or showed the opposite pattern (candidate substrates). This was based on the assumption that the levels of metabolic products and substrates of an enzyme correlate with the transcript levels of the corresponding gene. The peak areas of the features of interest were determined by manual integration in DataAnalysis and divided by the peak area of the manually integrated *m*/*z* trace of the [M+H]^+^ ion of the internal standard and the sample dry weight to calculate more accurate (i.e., peak area-based) FCs.

Metabolites were putatively and partially identified on the basis of ion types, accurate *m*/*z* values and intensities of parent and fragment ions. Sugar moieties were identified on the basis of accurate calculations of neutral losses; this allowed differentiation between a hexosyl (neutral loss: 162.0528 Da) and a caffeoyl (162.0317 Da) moiety as well as between a deoxyhexosyl (146.0579 Da) and a *p*-coumaroyl (146.0368 Da) moiety. Since deoxyhexosyl substitutions in flavonoids are most commonly rhamnosyl moieties, we assumed rhamnosyl groups as deoxyhexosyl substitutions/neutral losses. For structural formula prediction, *in-silico* fragmentation with MetFrag (Ruttkies *et al*., 2016) was applied to the ESI^+^ fragments, using the PubChem database (Kim *et al*., 2019); this was accompanied by spectral matching with entries in the MassBank of North America (https://mona.fiehnlab.ucdavis.edu/).

## Results and discussion

### Characterization of BGLU expression variant lines for functional studies

The reliability of the results of reverse genetic approaches strongly depends on the availability of suitable, thoroughly characterized mutants. The selected *bglu* T-DNA insertion mutants (*bglu1-1*, *bglu3-2*, *bglu4-2*) were characterized by long-read sequencing and additional Sanger sequencing of amplicons across both T-DNA::genome junctions. These characterizations demonstrated homozygosity of the T-DNA insertion alleles (Supplementary Fig. S1) and single T-DNA insertions in the corresponding *BGLU* genes, without loss of neighboring genes. They also revealed the exact positions, orientations and border sequences of the T-DNA insertions (Fig. 1A, Supplementary Table S2). Thus, the mutants were sufficiently characterized according to current standards (Pucker *et al*., 2021; Ülker *et al*., 2008). In *bglu1-1*, the T-DNA was inserted at pseudochromosome 1 position 17,116,579 [right T-DNA border (RB)] and position 17,116,598 [left T-DNA border (LB)] in the third intron of *BGLU1*. In *bglu3-2*, the insertion was located at pseudochromosome 4 positions 11,709,058 (LB) and 11,708,997 (RB), with the LB in the sixth exon and the RB in the sixth intron of *BGLU3*. In *bglu4-2*, the T-DNA was inserted at pseudochromosome 1 positions 22,156,000 (LB) and 22,156,017 (LB) in the third intron of *BGLU4*. The two LBs were probably derived from an insertion in LB-RB::RB-LB configuration (Kleinboelting *et al*., 2015; Pucker *et al*., 2021). In almost all cases, there were additional nucleotides at the T-DNA::genome junctions (Supplementary Table S2) as described for DNA double strand break-based T-DNA insertions (Kleinboelting *et al*., 2015).

**Fig. 1.**
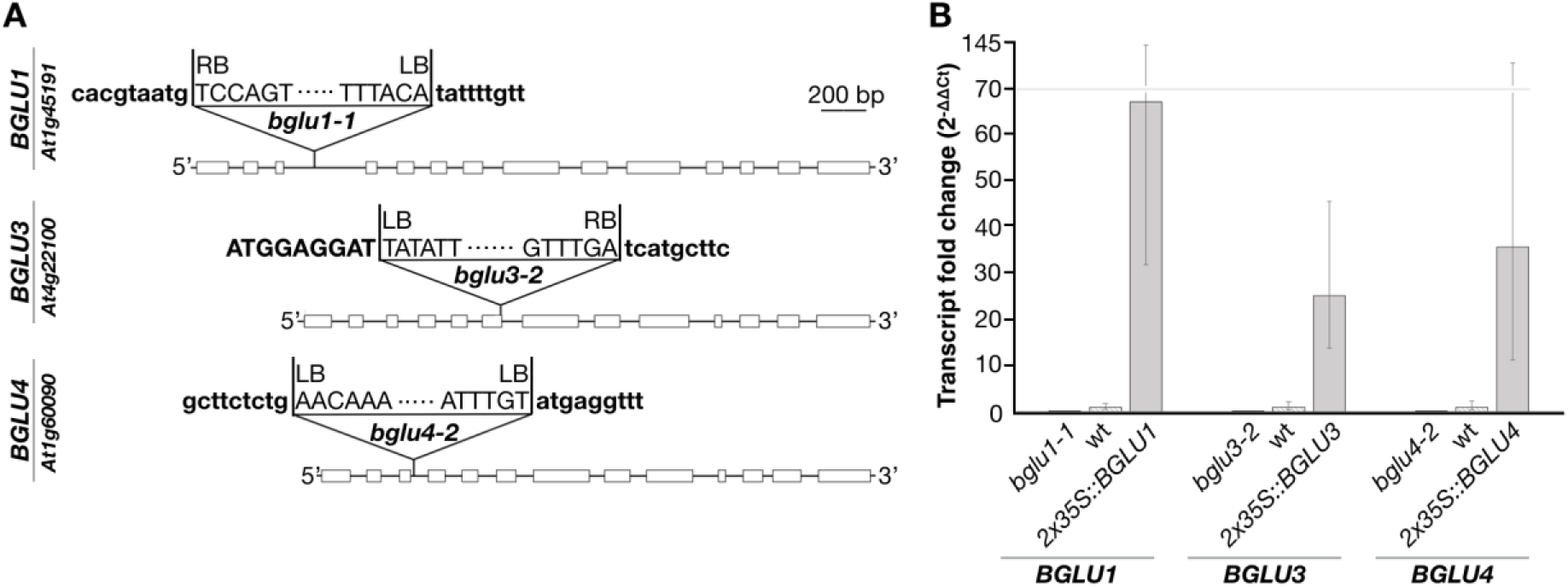
Characterization of *A. thaliana bglu* T-DNA insertion mutants and *2x35S::BGLU* overexpression lines for *BGLU1*, *BGLU3* and *BGLU4*. (A) *bglu* T-DNA insertion alleles with boxes indicating coding exons of the wild type (wt) allele structure, lines between the boxes indicating introns and triangles marking the positions of T-DNA insertions. T-DNA insertion boundaries are shown with the genomic sequences given in bold (introns in lowercase, exons in uppercase). Sequences that originate from the insertion event and do not match either genomic *BGLU* or T-DNA sequences are not shown here (for them, see Supplementary Table S2). (B) Differential *BGLU* gene expression of the intact transcripts of *BGLU1*, *BGLU3* and *BGLU4* in the *BGLU* expression variants, relative to the wt. Transcript levels were measured by RT-qPCR in rosette leaves (*BGLU1*), dry mature seeds (*BGLU3*) and 24 h water-soaked mature seeds (*BGLU4*), respectively. Data presented are from three biological replicates with three technical replicates each; error bars indicate the asymmetrically distributed cumulative standard error. The y-axis is compressed between 70 and 145.

With loss-of-function mutants, the functions of genes and the proteins encoded by them can be revealed, but functional redundancy of genes can prevent or mask a clear phenotype in single loss-of-function mutants (Bolle *et al*., 2013; O’Malley and Ecker, 2010). To overcome this limitation, we also included overexpression lines. The full-length CDSs of *BGLU1*, *BGLU3* and *BGLU4* under the control of the 2x35S promotor were stably introduced into *A. thaliana*, generating ten *2x35S::BGLU1*, three *2x35S::BGLU3* and ten *2x35S::BGLU4* overexpression lines. Transgene expression was confirmed in rosette leaves by RT-PCR (Supplementary Fig. S2). For each *BGLU* gene, the line with the strongest *BGLU* overexpression was selected for further experiments: *2x35S::BGLU1* #6, *2x35S::BGLU3* #44 and *2x35S::BGLU4* #91. For final characterization, intact *BGLU* transcripts were examined by RT-qPCR and compared between the genotypes. The plant parts with highest gene expression levels in the wt (see Material and Methods) were chosen: rosette leaves for *BGLU1*, dry mature seeds for *BGLU3* and 24 h water-soaked mature seeds for *BGLU4*. Compared to the wt, the insertion mutants showed lower (FC: 0.04 for *bglu1-1*, 0.001 for *bglu3-2*, 0.0005 for *bglu4-2*) and the overexpression lines much higher (FC: 67.3 for *2x35S::BGLU1*, 25.1 for *2x35S::BGLU3*, 35.8 for *2x35S::BGLU4*) relative target gene expression values (Fig. 1B). Thus, the chosen *BGLU* expression variant lines were suitable to screen for potential metabolic products and substrates. Several metabolic features possibly representing products and substrates of the encoded enzymes were found (see below; Fig. 2, Supplementary Table S3, S4, S5). Feature names are based on the RT and *m*/*z* values derived from feature picking and subsequent alignment across samples; *m*/*z* and RT values of the mass spectra may differ slightly from these values, because spectra were taken from single samples with high feature intensities.

**Fig. 2.**
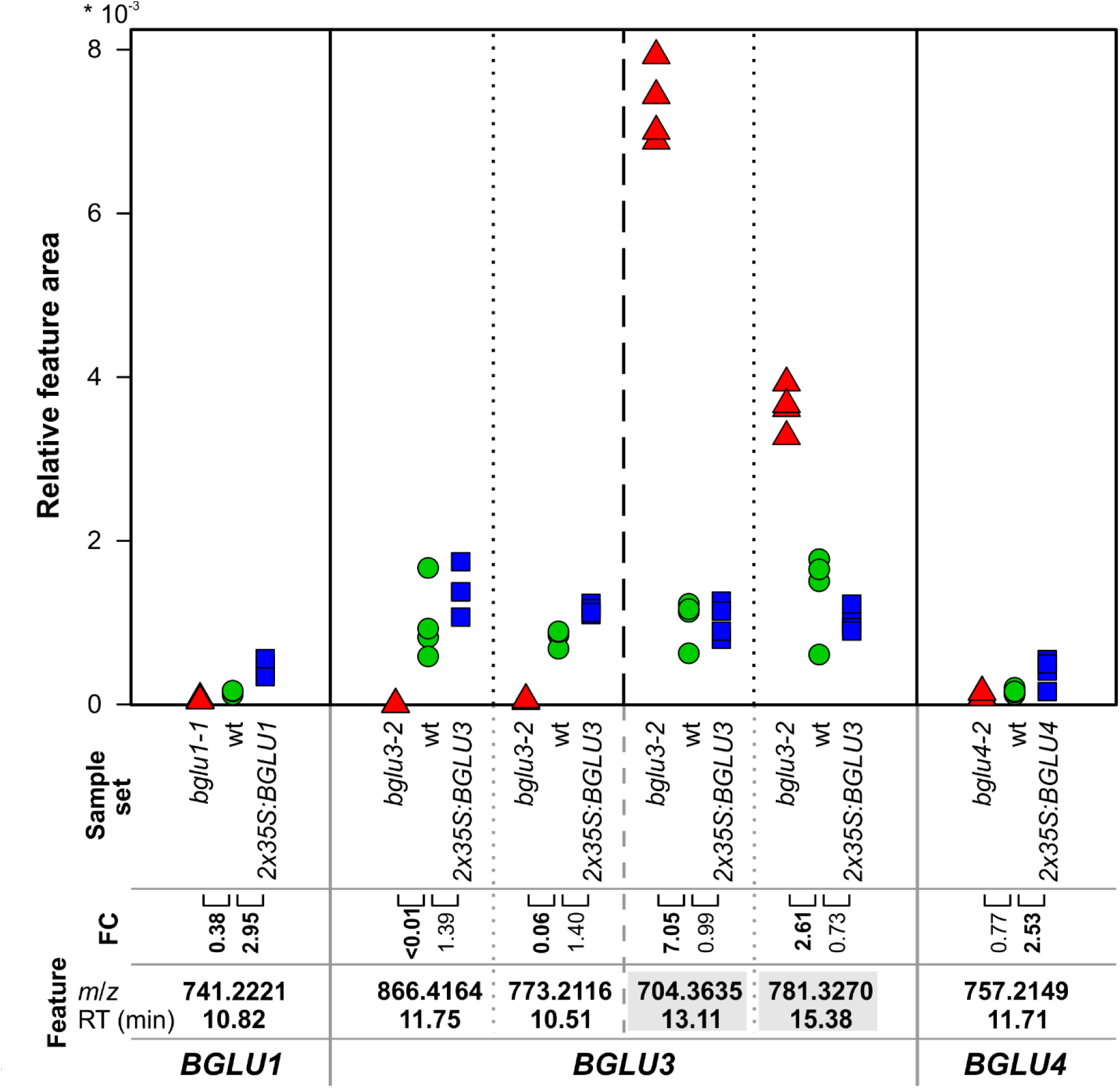
Main candidate product and substrate metabolic features of *A. thaliana BGLU1*, *BGLU3* and *BGLU4*. Feature intensities in rosette leaves (*BGLU1*), dry mature seeds (*BGLU3*) and 24 h water-soaked mature seeds (*BGLU4*), respectively, are given as relative feature areas, i.e., peak areas divided by the peak area of the internal standard and by the dry weight of the samples. The areas of the different features are not directly comparable, as different spectra rates were used. Peaks of *bglu* mutant samples are indicated by red triangles, those of wt samples by green circles and those of *2x35S::BGLU* samples by blue squares. Vertical solid lines separate the candidate features belonging to *BGLU1*, *BGLU3* and *BGLU4*; therein, candidate products (left) and substrates (right, feature names on gray background) are separated by vertical dashed lines and features within these groups by vertical dotted lines. Fold changes (FCs, in bold if ≥ 1.5 or ≤ 0.67) were calculated as the mean peak areas of the *BGLU* expression variants divided by the mean peak areas of the wt; n = 4 biological replicates.

### BGLU1 is a putative GH1-type flavonol glycosyltransferase

In the *BGLU1* data set (rosette leaves of wt, *bglu1-1*, *2x35S::BGLU1*), one candidate product feature (*m*/*z* value of 741.2221 at RT 10.82 min) was found, whose intensity pattern mirrored the gene expression pattern (Fig. 2, Supplementary Fig. S3). Low *BGLU1* transcript levels were still detected in the insertion mutant *bglu1-1*, probably due to the fact that the inserted T-DNA is located in the middle of a long intron (Fig. 1A) and that the primary transcript of the insertion allele was presumably spliced correctly in some cases. This may explain the occurrence of the candidate product feature (*m*/*z* of 741) at low levels in the *bglu1-1* samples.

There are several indications that the putative product of BGLU1 (feature with *m*/*z* 741.2221 at 10.82 min) is a triglycosylated kaempferol with the molecular formula C_33_H_40_O_19_ and an average molecular weight of 740.6606 Da. The presence of three *O*-bound sugars is suggested by the successive loss of two deoxyhexosyl (probably rhamnosyl) moieties (fragment with *m*/*z* 595: [precursor–146]^+^; fragment with *m*/*z* 449: [precursor–146–146]^+^) and one hexosyl moiety (fragment with *m*/*z* 287: [precursor–146–146–162]^+^) in the ESI^+^ mode (Fig. 3A). In addition, a minor fragment with an *m*/*z* value of 433 was visible, which can be explained by the loss of one rhamnosyl and one hexosyl moiety ([precursor–146–162]^+^).

**Fig. 3.**
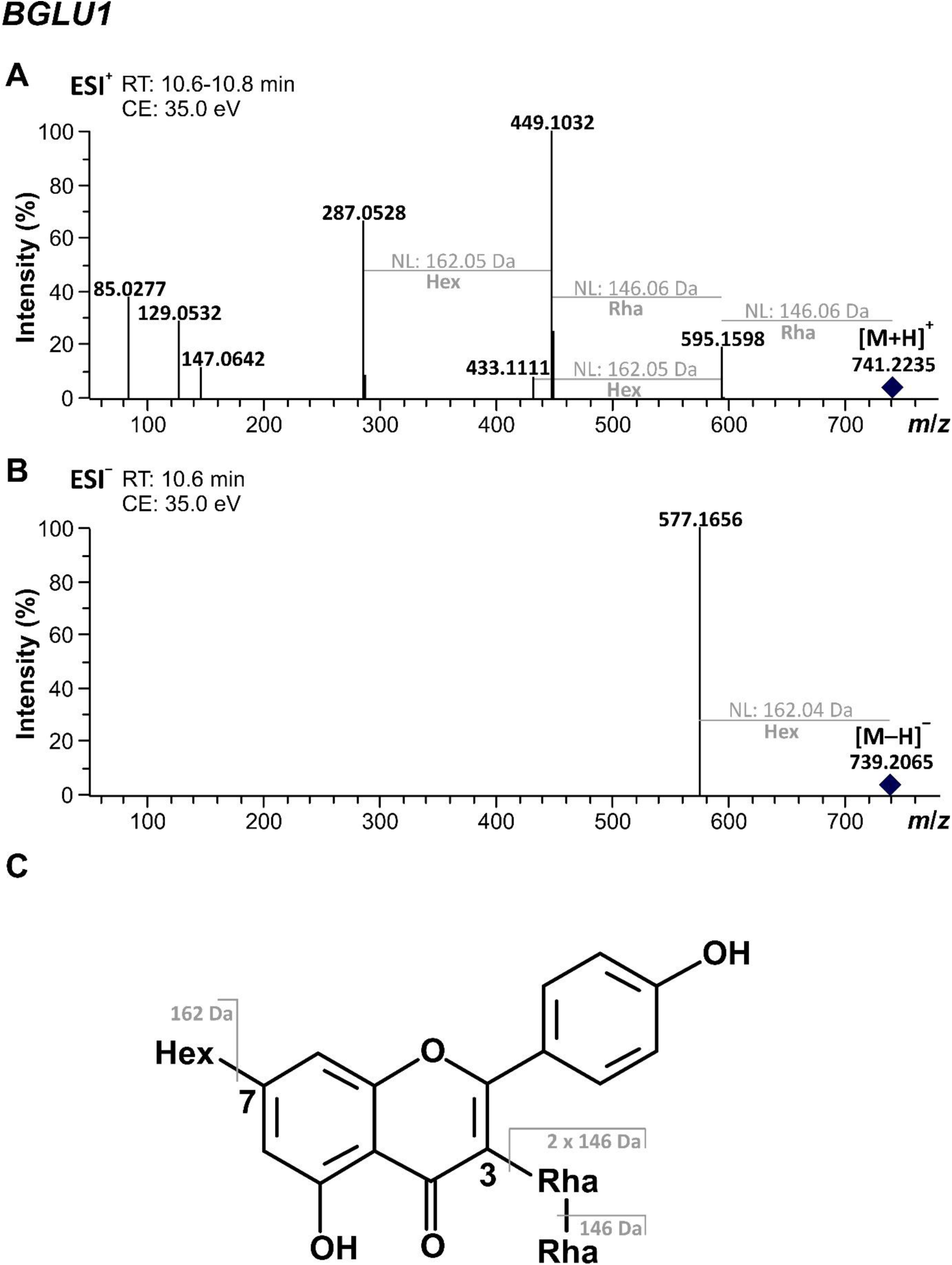
Candidate product of BGLU1 of *A. thaliana*. While (A) shows the MS/MS spectrum of the candidate product feature in the ESI^+^ mode, (B) shows the fragmentation of the corresponding ion found in ESI^−^ mode. The assumed ion types are given at the corresponding *m*/*z* values. Precursor ions are indicated by diamonds. (C) Assumed structure of the proposed metabolite and possible fragmentation pattern. All glycosyl groups are assumed to be *O*-bound. CE, collision energy; Hex, hexosyl; NL, neutral loss; Rha, rhamnosyl.

The fragment with an *m*/*z* value of 287 indicates that the precursor (parent ion *m*/*z* 741) may be an [M+H]^+^ ion with kaempferol (a flavonol) as aglycon or an [M]^+^ ion with cyanidin (an anthocyanidin) as aglycon. Glycosides with these backbones can be distinguished by the ion types they form in ESI^−^ mode (Sun *et al*., 2012). While flavonol glycosides produce mainly [M–H]^−^ ions, anthocyanins form pronounced [M– 2H]^−^ and [M–2H+H_2_O]^−^ ions as well as formic acid adducts related to both ion types; doubly charged ions corresponding to these ions also may occur. In the ESI^−^ mode, we detected an ion with an *m*/*z* value of 739, which may be an [M–H]^−^ ion of a kaempferol triglycoside or an [M–2H]^−^ ion of a cyanidin triglycoside. The occurrence of a putative formic acid adduct (*m*/*z* 785) of this ion at the same RT does not fit to a flavonol, as these usually do not show formic acid adducts. However, the facts that there was no obvious ion with *m*/*z* 757, which would be the [M–2H+H_2_O]^−^ ion expected for a cyanidin triglycoside and that there was no formic acid adduct of this ion suggest that the aglycon may be kaempferol. Fragmentation of the ion (*m*/*z* 739) revealed a dominant fragment with an *m*/*z* value of 577 (Fig. 3B), supporting that the metabolite contains an *O*-bound hexosyl moiety; the slight deviation of the accurate mass of the neutral loss (162.04) from the expected value (162.05) may be due to the low peak intensities. In contrast to the ESI^+^ mode, in ESI^−^ mode no fragments belonging to further losses of sugar moieties and no aglycon fragment were found, making further conclusions difficult. *In-silico* fragmentation in MetFrag using the ESI^+^ data further supported a triglycosylated kaempferol, as the two best hits were kaempferol 3-[6-*O*-(3-*O*-α-L-rhamnopyranosyl-α-L-rhamnopyranosyl)-β-D-glucopyranosyloxide] (PubChem CID: 101502553) and kaempferol 3-*O*-[α-L-rhamnopyranosyl(1→2)-β-D-galactopyranosyl]-7-*O*-α-L-rhamnopyranoside) (CID: 57397583).

Our putative metabolite identification is in good agreement with published results (Wu *et al*., 2018). Genome-wide association studies, using untargeted LC-MS metabolic fingerprinting of 309 *A. thaliana* accessions, revealed a feature (*m*/*z* 741.2220, ESI^+^) with a MS/MS spectrum similar to the one we found. This metabolic trait was traced back to the *BGLU1* locus. The metabolite-gene correlation was validated by the analysis of a *bglu1* mutant (SALK_060948 allele), which showed lower levels of the feature with *m*/*z* 741.2220. The authors were unable to identify the compound but showed that the feature was also associated with the *UGT78D1* locus, which encodes a kaempferol and quercetin 3-*O*-rhamnosyltransferase UGT (Jones *et al*., 2003), suggesting that the metabolite may be 3-*O*-rhamnosylated.

In general, diverse kaempferol *O*-glycosides occur in *A. thaliana* leaves, varying in the number, type and positions of sugars (Hectors *et al*., 2014; Tohge *et al*., 2005), with C-3 and C-7 being the most common glycosylation sites (Stobiecki *et al*., 2006). For the feature of interest in the current study (*m*/*z* 741), the assumed loss of a hexosyl moiety from the [M–H]^−^ ion observed in ESI^−^ mode (fragment with *m*/*z* 577) indicates that the hexosyl group is bound at a terminal position. The C-3 position is more prone to fragmentation in ESI^+^ mode, whereas the C-7 position is readily fragmented in ESI^−^ mode (Kachlicki *et al*., 2016; Stobiecki *et al*., 2006). Thus, the prominent fragment with *m*/*z* 449 in ESI^+^ mode ([M+H–146–146]^+^) suggests that a rhamnosyl-rhamnosyl moiety is bound to the C-3 position, whereas the dominant fragment with *m*/*z* 577 in ESI^−^ mode ([M–H–162]^−^) may indicate that the hexosyl moiety is linked to the C-7 position.

Taken together, although we cannot rule out that the sugars are attached at other positions, our study provides evidence that the feature (*m*/*z* 741.2221 at 10.82 min) is a kaempferol 3-*O*-dirhamnoside 7-*O*-hexoside. We hypothesize that BGLU1 is a GH1-type flavonol glycosyltransferase catalyzing the transfer of a further sugar to an already di-glycosylated kaempferol derivative with a rhamnosyl moiety at the C-3 position (see hints from literature above). BGLU1 may transfer either a hexosyl moiety to the C-7 position of kaempferol 3-*O*-dirhamnoside or a second rhamnosyl to the C-3 rhamnosyl moiety of kaempferol 3-*O*-rhamnoside 7-*O*-hexoside. Phylogenetic clustering of GTs correlates with the glycosylation sites of the substrates (Jackson *et al*., 2011; Vogt and Jones, 2000) and BGLU1 clusters with BGLU6 (Ishihara *et al*., 2016; Miyahara *et al*., 2011). BGLU6 catalyzes the transfer of another glucosyl moiety to the glucosyl moiety at C-3 of a flavonol 3-*O*-glucoside 7-*O*-rhamnoside in *A. thaliana* (Ishihara *et al*., 2016). Therefore, we assume that BGLU1 is responsible for the transfer of a second rhamnosyl to the rhamnosyl moiety at C-3. However, to our knowledge, GH1-type flavonol rhamnosyltransferases have not been reported in plants so far. Since no candidate substrate feature was found for BGLU1, we cannot exclude that BGLU1 catalyzes the sugar transfer to a monoglycoside (presumably kaempferol 3-*O*-rhamnoside, see above) and that the third sugar is transferred by another enzyme; in this case, the intermediate diglycoside might have been below the detection limit, perhaps due to rapid metabolism. We also cannot fully rule out a *β*-glycosidase activity of BGLU1, but there were no candidate features indicating this.

### BGLU3 is a putative multifunctional GH1-type flavonoid glycosyltransferase

For the BGLU3 data set (dry mature seeds of wt, *bglu3-2*, *2x35S::BGLU3*), the FC screening revealed three main candidate product features and three main candidate substrate features, each with high peak intensities. In addition, some minor or less intense candidate features were detected (Fig. 2, Supplementary Table S3, S5). The intensity patterns of these features were generally consistent with the *BGLU3* transcript levels (Fig. 2, Supplementary Fig. S3, Supplementary Table S5). Relatively similar feature intensities were determined in the wt and *2x35S::BGLU3* samples, but lower (product features) and higher (substrate features) intensities, respectively, in the *bglu3-2* mutant. The reason for the comparable transcript levels in *2x35S::BGLU3* and wt is unclear. Although insertion of the transgene into a region with low transcriptional activity (Nagaya *et al*., 2005) cannot be ruled out, metabolite-mediated negative feedback regulation of gene expression (Xu *et al*., 1999) might also be involved. The occurrence of candidate product features at low levels in the *bglu3-2* mutant, which lacks *BGLU3* expression, may indicate functional redundancies conferred by other enzymes.

The candidate product feature with an *m*/*z* value of 866.4164 at 11.75 min and the corresponding putative substrate (*m*/*z* 704.3635 at 13.11 min) indicate a hexosyltransferase activity of BGLU3, as the *m*/*z* difference between these ions of 162.05 matches to a hexosyl moiety. The product-substrate relationship between these features is further supported by their similar fragmentation patterns (Fig. 4A, 4B). Both features showed a major fragment indicating a CO_2_ loss ([precursor–44]^+^; fragments with *m*/*z* 822 and *m*/*z* 660 for the candidate product and candidate substrate feature, respectively), which may indicate a carboxyl function in the metabolites. Furthermore, a neutral loss of 226 Da (fragments with *m*/*z* 640 and *m*/*z* 478, respectively) and fragments at *m*/*z* values of 398 and 339 were observed for both. In addition to the similar fragmentation, both features showed co-eluting doubly charged features. While the feature with *m*/*z* 433.7128 at 11.76 min probably represents the doubly-charged version ([M+H]^2+^, explanation see below) of the candidate product feature (*m*/*z* 866), the feature with *m*/*z* 330.6915 at 13.11 min may be an [M+H-CO_2_]^2+^ ion belonging to the candidate substrate feature (*m*/*z* 704) (Supplementary Table S3). The structure of the metabolites could not be determined. However, the high *m*/*z* may indicate that they are condensed flavonoids. The doubly charged ions may be due to the presence of a naturally positively charged anthocyani(di)n moiety ([M]^+^ ion), which together with a protonation at another unit of the metabolite leads to a double charge ([M+H]^2+^). Although some fragments pointed to a carboxypyranomalvidin-hexoside (a pyranoanthocyanin) (Fulcrand *et al*., 1998) and an (epi)gallocatechin (a flavanol) (Nave *et al*., 2010; Sánchez-Ilárduya *et al*., 2012) as potential units of the candidate product, the overall *m*/*z* of the candidate product and substrate features did not support this combination. Some further minor candidate product features were found, many of them being doubly charged as well and showing mixtures of singly and doubly charged fragments (Supplementary Table S3, S5). Some of the corresponding metabolites may be derived from the metabolite belonging to the feature with *m*/*z* 866 via further biosynthesis pathways (Supplementary Table S4). Among the minor candidate features, the product with *m*/*z* 411.7178 (11.87 min; doubly charged, putatively [M+H]^2+^) may be related to the substrate with *m*/*z* 660.3742 (13.21 min; singly charged, putatively [M]^+^), with the features differing in a hexosyl moiety. This product/substrate pair seems to be chemically similar to the product/substrate pair with *m*/*z* 866 (433 in doubly charged version) / 704, with a carboxyl function less (44 Da difference, CO_2_).

**Fig. 4.**
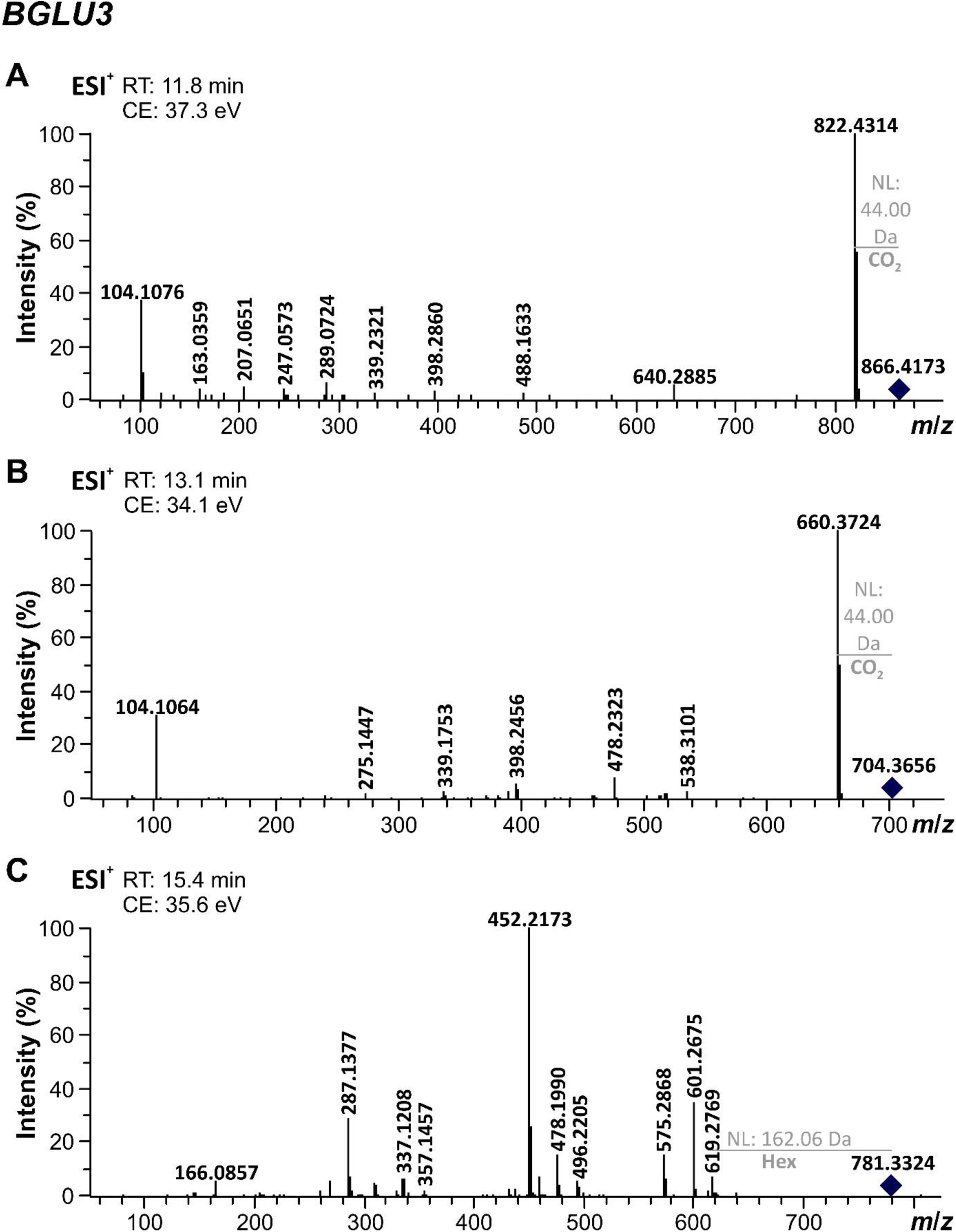
Main candidate product and substrates of BGLU3 of *A. thaliana*. MS/MS spectra (ESI^+^ mode) of (A) candidate product feature with an *m*/*z* value of 866.4164 at a retention time (RT) of 11.75 min, (B) candidate substrate feature with an *m*/*z* value of 704.3635 at 13.11 min, (C) candidate substrate feature with an *m*/*z* value of 781.3270 at 15.38 min. Precursor ions are indicated by diamonds. CE, collision energy; NL, neutral loss.

The structure of the candidate substrate feature with *m*/*z* 781.3270 at 15.38 min could also not be resolved. However, this substrate seems to contain a hexosyl moiety (neutral loss of 162.06 Da, fragment *m*/*z* 619) and we found a minor candidate product feature with *m*/*z* 943.3804 at 13.74 min, which differed by 162.05 Da (i.e., a hexosyl) and showed some similar MS/MS fragments (Fig. 4C, Supplementary Tables S3, S5). This suggests that BGLU3 transfers an additional hexosyl moiety. Some of the minor candidate BGLU3 substrate features showed similar characteristics and thus probably represent similar metabolites (Supplementary Table S3). More studies on flavonoids and their condensation reactions are needed to provide further information on whether the features observed in our study are (condensed) flavonoids. In any case, our study suggests that BGLU3 acts in the seeds and transfers hexosyl moieties to complex substrates.

Our results provide some evidence that BGLU3 also catalyzes a reaction leading to a triglycosylated quercetin with the molecular formula C_33_H_40_O_21_ and an average molecular weight of 772.6594 Da, represented by the candidate product feature with an *m*/*z* of 773.2116 at 10.51 min (Fig. 2, Supplementary Tables S3, S5). The MS/MS fragments with *m*/*z* 449 ([precursor–324]^+^) and *m*/*z* 303 ([precursor–324–146]^+^) indicate the loss of two *O*-bound hexosyl moieties and of one *O*-bound deoxyhexosyl (likely rhamnosyl) moiety (Fig. 5A). Based on the fragment with *m*/*z* 303, the parent ion (*m*/*z* 773) could be either an [M+H]^+^ ion with quercetin (a flavonol) as aglycon or an [M]^+^ ion with delphinidin (an anthocyanidin) as aglycon. In the ESI^-^ mode, the ion with *m*/*z* 771 (Fig. 5B) could represent the [M–H]^−^ ion of a quercetin triglycoside or the [M–2H]^−^ ion of a delphinidin triglycoside; as no obvious [M–2H+H_2_O]^−^ ion (*m*/*z* 789) and no formic acid adducts were found, which would indicate an anthocyanidin aglycon (Sun *et al*., 2012), we assume that the aglycon is quercetin. Fragmentation of the ion with *m*/*z* 771 supports the presence of one *O*-bound rhamnosyl ([M–H– 146]^−^; fragment *m*/*z* 625) and two *O*-bound hexosyl moieties ([M–H–324]^−^; *m*/*z* 447) (Fig. 5B). However, as no aglycon fragment was found in ESI^-^ mode, potentially due to non-optimal collision energies, the identity of the aglycon cannot be further assessed. Nevertheless, *in-silico* fragmentation using the ESI^+^ data also revealed quercetin triglycosides as best hits. Various quercetin *O*-glycosides that differ in the number, type and positions (mainly C-3 and C-7) of the sugars occur in *A. thaliana* seeds (Kerhoas *et al*., 2006; Lepiniec *et al*., 2006; Routaboul *et al*., 2012; Saito *et al*., 2013). The neutral loss of a rhamnosyl in ESI^−^ mode (fragment with *m*/*z* 625) from the [M–H]^−^ ion suggests a terminal position of the rhamnosyl moiety. This and the loss of a hexosylhexosyl moiety from the parent ions ([M+H–324]^+^, fragment *m*/*z* 449; [M–H–324]^−^, fragment *m*/*z* 447) may indicate the attachment of the rhamnosyl and hexosylhexosyl moieties at different positions. We argue that due to the loss of a hexosylhexosyl from the parent ion in ESI^+^ mode (fragment *m*/*z* 449), most likely from the fragmentation at C-3 (Kachlicki *et al*., 2016; Stobiecki *et al*., 2006), the feature with *m*/*z* 773 may represent a quercetin 3-*O*-dihexosyl 7-*O*-rhamnoside. However, we cannot rule out that the sugars are attached in a different way. Based on the assumption that the phylogenetic clustering of glycosyltransferases correlates with the glycosylation site (Jackson *et al*., 2011; Vogt and Jones, 2000) and on the clustering of BGLU3 with BGLU6 (Ishihara *et al*., 2016; Miyahara *et al*., 2011), BGLU3 may be responsible for the transfer of the second hexosyl moiety to the C-3 position. Nevertheless, other glycosylation reactions and a *β*-glycosidase activity cannot be excluded for BGLU3, for the same reasons as discussed for BGLU1.

**Fig. 5.**
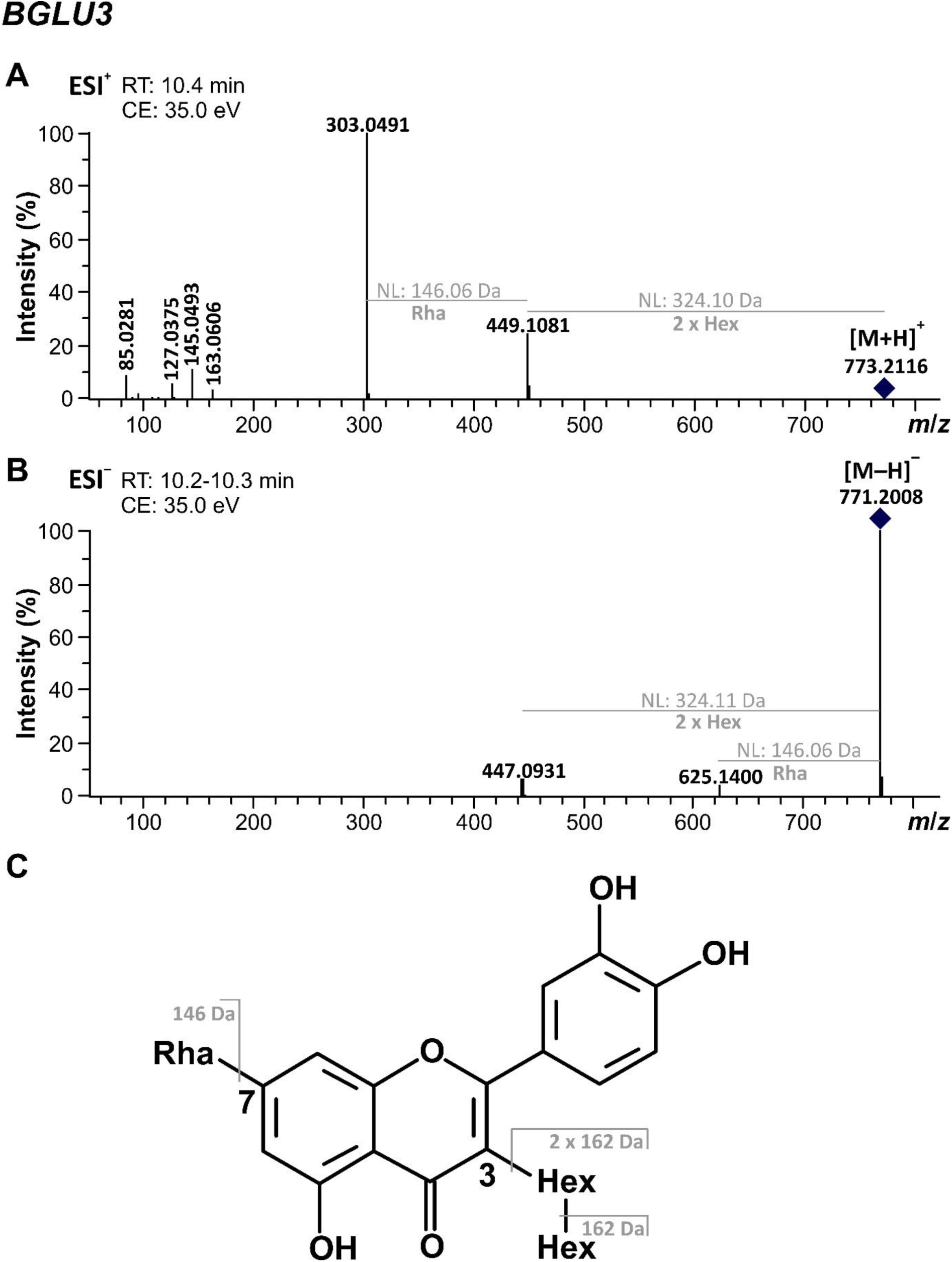
Candidate product of BGLU3 of *A. thaliana*. While (A) shows the MS/MS spectrum of the candidate product feature in the ESI^+^ mode, (B) shows the fragmentation of the corresponding ion found in ESI^−^ mode. The assumed ion types are given at the corresponding *m*/*z* values. Precursor ions are indicated by diamonds. (C) Assumed structure of the proposed metabolite and possible fragmentation pattern. All glycosyl groups are assumed to be *O*-bound. CE, collision energy; Hex, hexosyl; NL, neutral loss; Rha, rhamnosyl.

Taken together, our results indicate that BGLU3 acts as a GH1-type hexosyltransferase, transferring hexosyl moieties to different flavonoids. Multifunctionality is also known from the GH1-type GT Os9BGlu31 from rice, which transfers sugar moieties to compounds as diverse as flavonoids, phenolic acids and phytohormones (Luang *et al*., 2013). Based on RNA-Seq data (TraVa database), BGLU3 expression is restricted to seeds, suggesting that the proposed glycosylation reactions are specific to these plant parts. Indeed, seeds are known to contain various flavonoids (Kerhoas *et al*., 2006), some of them being exclusively found in mature seeds as shown for flavonol 3-*O*-sophoroside 7-*O*-rhamnosides in tomato seeds (Alseekh *et al*., 2020) or epicatechins in *A. thaliana* seeds (Saito *et al*., 2013).

### BGLU4 is a putative GH1-type flavonol glycosyltransferase

The FC screening in the BGLU4 data set (24 h water-soaked mature seeds of wt, *bglu4-2*, *2x35S::BGLU4*) revealed one candidate product feature (*m*/*z* of 757.2149 at 11.71 min; Fig. 2). The pattern of feature intensities was largely similar to that of *BGLU4* gene expression (Fig. 2, Supplementary Fig. S3). However, the candidate product was also found in traces in the *bglu4-2* mutant, which showed no *BGLU4* expression, suggesting functional redundancy.

The metabolic analyses suggest that the BGLU4 candidate product (*m*/*z* 757.2149 at 11.71 min) could be a triglycosylated quercetin (molecular formula C_33_H_40_O_20_, average molecular weight: 756.6600 Da). The fragmentation of this feature in the ESI^+^ mode (Fig. 6A) suggests a loss of a rhamnosyl and a hexosyl moiety (fragment *m*/*z* 449, [precursor–146–162]^+^) and a further loss of a rhamnosyl moiety (fragment *m*/*z* 303; [precursor–146–162–146]^+^). The fragment with *m*/*z* 303 indicates that the parent ion (*m*/*z* 757) could be either an [M+H]^+^ ion with quercetin as aglycon or an [M]^+^ ion with delphinidin as aglycon. Measurements in the ESI^−^ mode (Fig. 6B) revealed an ion with *m*/*z* 755, which could be the [M–H]^−^ ion of a quercetin triglycoside or the [M–2H]^−^ ion of a delphinidin triglycoside. We suggest that the aglycon is a quercetin, because no obvious ion with *m*/*z* 773 ([M–2H+H_2_O]^−^ and no formic acid adducts were found, which would be expected for a delphinidin triglycoside (Sun *et al*., 2012), and because *A. thaliana* lacks a gene encoding F3’5’H activity (Falginella *et al*., 2010). In addition, the ESI^−^ data support the existence of three sugar moieties (fragment *m*/*z* 609: [precursor–146]^−^; *m*/*z* 447: [precursor–146– 162]^−^; *m*/*z* 301: [precursor–146–162–146]^−^), although the masses of the neutral losses deviate somewhat, probably due to low intensities. Furthermore, *in-silico* fragmentation with the ESI^+^ data revealed glycosylated quercetins as best hits, most likely quercetin 3-*O*-[α-L-rhamnopyranosyl(1→2)-β-D-galactopyranosyl]-7-*O*-α-L-rhamnopyranoside (CID: 57397680), quercetin 3-rhamnosyl-(1→4)-rhamnosyl-(1→6)-glucoside (CID: 44259158) or quercetin 3-rhamnosyl-(1→2)-rhamnosyl-(1→6)-glucoside (CID: 44259157). In the ESI^+^ mode, the neutral loss of 308 Da (one rhamnosyl and one hexosyl), leading to the prominent fragment with *m*/*z* 449, is probably derived from fragmentation at C-3, whereas the loss of a rhamnosyl moiety (146 Da; fragment *m*/*z* 609) in the ESI^−^ mode indicates that this moiety is probably derived from C-7 (Kachlicki *et al*., 2016; Stobiecki *et al*., 2006). Based on the fragment with *m*/*z* 447 (ESI^−^) indicating a hexosyl loss from the fragment with *m*/*z* 609 (probably derived from the loss of a rhamnosyl at C-7, see above), we argue that at the C-3 position the rhamnosyl moiety is bound to the backbone and that the hexosyl moiety is bound to this rhamnosyl and located at the outer position. Thus, the product of BGLU4 may be a quercetin 3-*O*-rhamnosyl hexoside 7-*O*-rhamnoside. As discussed for the other BGLU proteins above, BGLU4 may transfer a hexosyl moiety to the already rhamnosylated C-3 position, while we cannot exclude the catalyzation of another sugar transfer or a *β*-glycosidase activity.

**Fig. 6.**
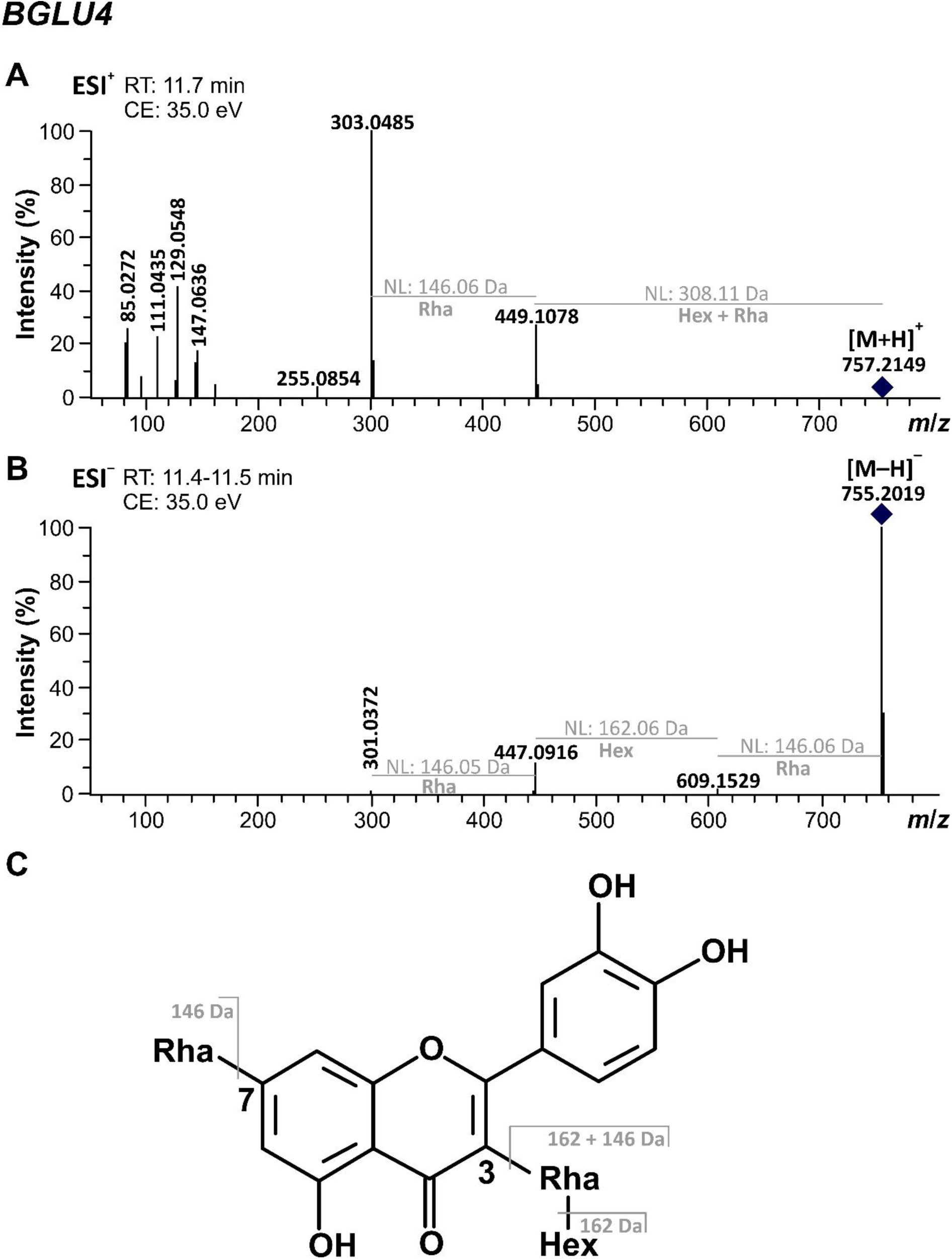
Candidate product of BGLU4 of *A. thaliana*. While (A) shows the MS/MS spectrum of the candidate product feature in the ESI^+^ mode, (B) shows the fragmentation of the corresponding ion found in ESI^−^ mode. The assumed ion types are given at the corresponding *m*/*z* values. Precursor ions are indicated by diamonds. (C) Assumed structure of the proposed metabolite and possible fragmentation pattern. All glycosyl groups are assumed to be *O*-bound. CE, collision energy; Hex, hexosyl; NL, neutral loss; Rha, rhamnosyl.

### BGLU proteins are localized in different cell compartments

GH1-type GTs are assumed to specifically catalyze the glycosylation of already substituted flavonoids, taking place either at the backbone, at another sugar moiety or at an acyl moiety (Ishihara *et al*., 2016; Matsuba *et al*., 2010; Miyahara *et al*., 2013). Most of these enzymes have a predicted localization signal for the vacuole, where flavonoid modifications by GHs and GTs also seem to take place (Luang *et al*., 2013; Sasaki *et al*., 2014). In subcellular localization experiments using BY-2 protoplasts, all BGLU-RFP fusion proteins were found to be localized in the cytoplasm, mostly at endomembranes around the nucleus (Fig. 7). BGLU3 and BGLU4 were also localized in the nucleus. In addition, all proteins (especially BGLU4) showed a weak localization in the vacuole in some protoplasts. Likewise, Ishihara *et al*. (2016) revealed subcellular localization of BGLU6 in the cytoplasm and Matsuba *et al*. (2010) demonstrated that one AGT is localized in the cytoplasm and vacuole. In the present study, the subcellular endomembrane control fusion protein TT13-GFP showed co-localization of the investigated BGLUs with endomembranes around the nucleus, probably the rough endoplasmatic reticulum (ER) (Fig. 7). This suggests that the BGLU proteins are synthesized at the rough ER and are subsequently transported to the vacuole by vesicular traffic through the cytoplasm.

**Fig. 7.**
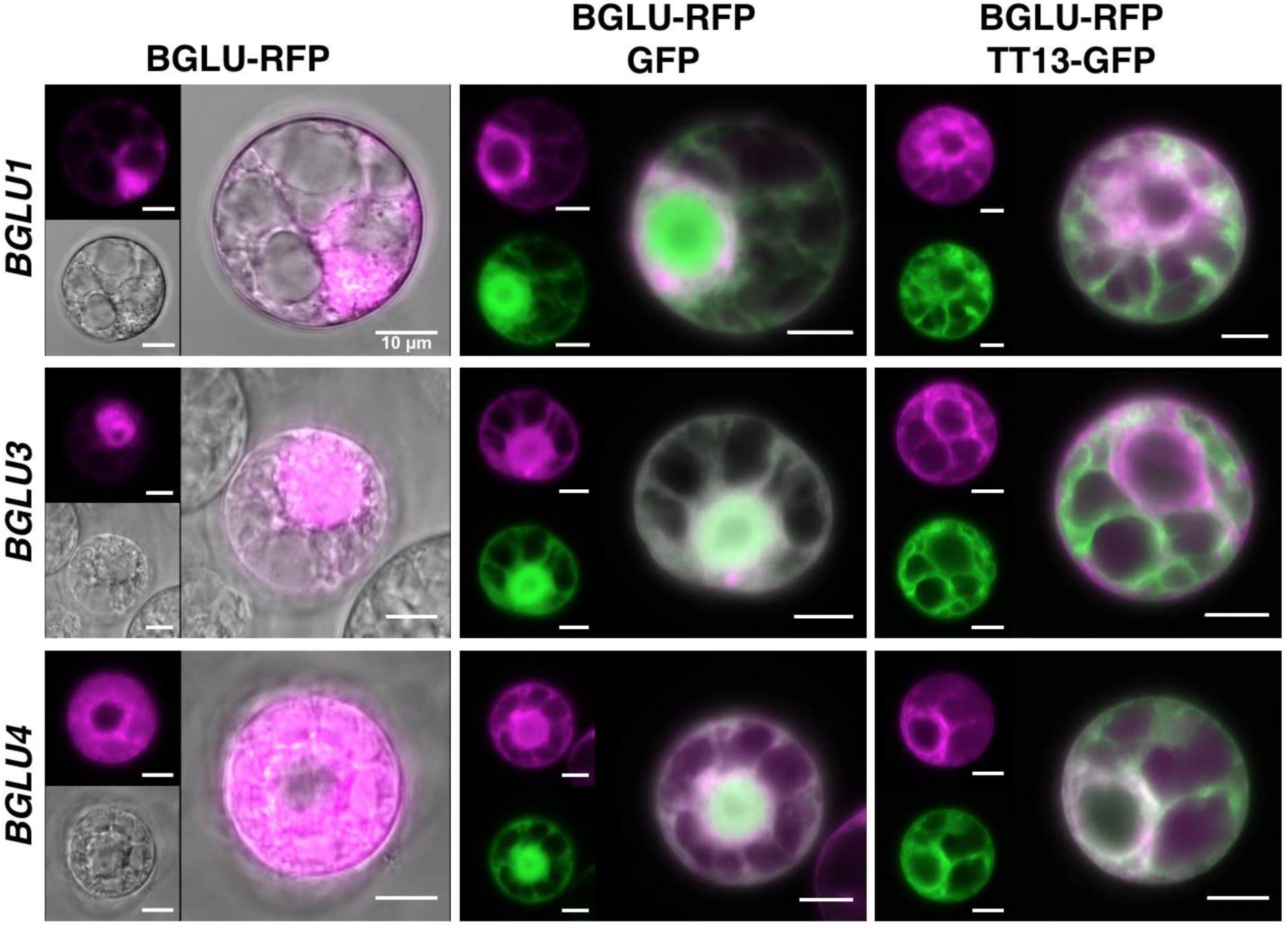
Subcellular localization of BGLU1, BGLU3 and BGLU4. Pictures show transient accumulation of BGLU-RFP fusion proteins in BY2 protoplasts. The second and third columns show co-localization with GFP (nucleus and cytoplasm) and TT13-GFP (vacuole), respectively. Co-localizations with positive signals for GFP and RFP in the merged images are visualized in white. Scale bars: 10 µm.

### Putative functions and relevance of GH1-type GTs

In summary, the screening of metabolites in different *BGLU* genotypes with genetically modified expression levels by metabolic fingerprinting suggests that the proteins encoded by *BGLU1*, *BGLU3* and *BGLU4* possess GH1-type glycosylation activity on glycosylated flavonoids. All three BGLUs appear to be involved in the synthesis of triglycosides, with BGLU1 most likely acting on a kaempferol diglycoside and BGLU3 and BGLU4 acting on different quercetin diglycosides. BGLU3 possibly also acts on other substrates, possibly condensed flavonoids.

Thus, our study suggests a broader range of functions of GH1-type GTs in plants than uncovered yet. The precise functions in terms of metabolic steps catalyzed by the enzymes and the biological functions of the metabolites remain to be determined. In general, it is suggested that GH1-type GTs act on already substituted flavonoids in the vacuole, where the acyl-sugar donor is located, while UGTs act in the cytoplasm, where they are co-localized with the UDP-sugar donor (Matsuba *et al*., 2010; Sasaki *et al*., 2014). Further studies on the subcellular localization and the activity of the enzymes in these compartments are needed. Also, the localization at endomembranes around the nucleus needs to be further clarified. If BGLU1, BGLU3 and BGLU4 are localized in the vacuole, they could act as AGTs, using sinapoyl-sugars as acyl-sugar donors, which are common in *A. thaliana* (Meißner *et al*., 2008; Miyahara *et al*., 2013).

The occurrence of many different GH1-type GTs may be related to the assumption that larger gene families, like the *BGLUs,* are based on plant part-specific expression of different paralogous genes (Cao *et al*., 2017; Gómez-Anduro *et al*., 2011). The glycosylated flavonoid products may be involved in plant responses to abiotic and biotic stresses, contributing to the enormous diversity of highly substituted flavonoids (Saito *et al*., 2013). Investigation of the transcriptional regulation of GH1-type GTs could provide insight into the biosynthetic pathways in which the enzymes are involved and their specific functions. Interestingly, Geng *et al*. (2021) were able to predict a gene co-expression module involved in flavonoid biosynthesis and stress response, including BGLU1, using the transcription factor activity-based expression prediction tool EXPLICIT. This module is based on transcription factors known to regulate structural flavonoid biosynthesis pathway genes.

Because the putative metabolite products were present at low levels in *bglu* mutants, future studies including multiple loss-of-function mutants should address potential functional redundancies and evolutionary sub-/neofunctionalization of *BGLU* genes. For example, such a functional redundancy is known for the *A. thaliana* anthranilate GTs UGT74F1 and UGT74F2 (Quiel and Bender, 2003) and is proposed for the *A. thaliana* AGTs BGLU10 and BGLU9 (Miyahara *et al*., 2013). Based on amino acid similarities (Miyahara *et al*., 2011) and gene expression in the same plant parts (TraVA, present study), BGLU5 and BGLU1 may have redundant roles in rosette leaves and the same may be the case for BGLU3 and BGLU4 in seeds. That BGLU3 may have a redundant enzymatic function with BGLU4 is supported by a minor candidate product feature (*m*/*z* 757.2166, 11.42 min) of BGLU3, exhibiting a similar fragmentation pattern as the BGLU4 candidate product with *m*/*z* 757.2149 at 11.71 min (Fig. 6; Supplementary Table S3), thus probably being an isomer of this compound. Finally, unambiguous structure validation requires measurements of reference standards in combination with NMR measurements. Both is challenging, because standards are not available for many complex flavonoids and NMR analyses require large amounts of purified compounds, which are almost impossible to generate, especially for metabolites from tiny *A. thaliana* seeds.

This work is a further step towards more detailed information about the abundance and function of GH1-type glycosyltransferases in plants and more precisely in *A. thaliana*. The data obtained also provide several hints towards the glycosylation activity on diverse substrates, which to our knowledge have not been reported in *A. thaliana* and are an interesting prospect for future studies.

## Abbreviations

2x35S: double enhancer cauliflower mosaic virus 35S
AGT: acyl-glucose-dependent glycosyltransferase
BGLU: *β*-glucosidase
CDS: coding sequence
CE: collision energy
Col-0: Columbia-0
ESI: electrospray ionization
FC: fold change
GH1: glycoside hydrolase family 1
GT: glycosyltransferase
Hex: hexosyl
MS/MS: spectrum, fragment spectrum
*m*/*z*: mass-to-charge ratio
NL: neutral loss
RT-qPCR: reverse transcriptase quantitative real-time PCR
rH: relative humidity
Rha: rhamnosyl
RT: retention time
RT-PCR: reverse transcriptase semi-quantitative PCR
T-DNA: transfer DNA
TT13: TRANSPARENT TESTA13
UGT: UDP-glycosyltransferase; wt, wild type

## Acknowledgements

We are grateful to Melanie Kuhlmann for her excellent assistance in the laboratory and to Andrea Voigt for her competent help in the greenhouse. We thank Hanna Marie Schilbert for providing her python script to assist RT-qPCR primer design.

## Funding

This work was supported by basic funding of the departments of Genetics and Genomics of Plants and Chemical Ecology provided by Bielefeld University/Faculty of Biology.

## Author contributions

RSt and BW conceived and designed the research. JFF, RSch and CM designed the metabolomics analyses. JFF, RSch, LMS and BP conducted experiments. BP generated, analyzed and deposited ONT data. RSch deposited the metabolic data in MetaboLights. JFF, RSt and RSch interpreted the data. JFF and RSch wrote the initial draft. RSt, CM and BW revised the manuscript. All authors read and approved the final manuscript.

## Data availability statement

ONT sequencing data were submitted to the European Nucleotide Archive under study ID PRJEB36305: bglu1-1 (ERS4255859), bglu3-2 (ERS4255860) and bglu4-2 (ERS4255861). Metabolic data (selected MS/MS spectra) will be made available upon publication of the paper in the MetaboLights repository (Haug *et al*., 2020; Haug *et al*., 2013) under the accession number MTBLS1965 (www.ebi.ac.uk/metabolights/MTBLS1965).

## Notes

### Competing Interest Statement

The authors have declared no competing interest.

