## Supplementary Tables S1-S4, Figures S1-S3 for "Metabolic fingerprinting reveals roles of *Arabidopsis thaliana* BGLU1, BGLU3 and BGLU4 in glycosylation of various flavonoids": Frommann_et_al_Supplement_BioRxiv.pdf

**Supplementary material**

**Table S1. Primers used in this study.**

| Usage | Target gene (allele) | Name | Sequence (5' → 3') |
|---|---|---|---|
| genotyping | *BGLU1* | RS1302<br>RS1303 | ACAGCAGGAGCGATTTCCCGGAAG<br>TGTAAGCCAGTTTCGGCCATGAG |
| | *bglu1-1* | RS1848<br>RS1845 | ATATTGACCATCATACTCATTGC<br>GGGACTTACCATGTTTGACAAGTT |
| | *BGLU3* | RS1319<br>RS1320 | GAGATGCAGCGACAAGAACGACTTCC<br>AGTTCCCTGATGAGCAGTTTCTACC |
| | *bglu3-2* | RS1846<br>RS1847 | ATAATAACGCTGCGGACATCTACATTTT<br>TTATTCACCAAGCAAACTCTTGAA |
| | *BGLU4* | RS1259<br>RS1260 | ATGGAACAGATTTTGGCTCTGTTTGCCATTTT<br>CTAGAAGGAAGAAGAGTATTTGCTCTGCA |
| | *bglu4-2* | RS1259<br>3144 | ATGGAACAGATTTTGGCTCTGTTTGCCATTTT<br>GTGGATTGATGTGATATCTCC |
| T-DNA border verification | *bglu1-1* | RS1302<br>F067 | ACAGCAGGAGCGATTTCCCGGAAG<br>CAGCAGAGCGCAGATACCAAATACTG |
| | *bglu1-1* | RS1845<br>RS1846 | GGGACTTACCATGTTTGACAAGTT<br>ATAATAACGCTGCGGACATCTACATTTT |
| | *bglu3-2* | 3144<br>RS1320 | GTGGATTGATGTGATATCTCC<br>AGTTCCCTGATGAGCAGTTTCTACC |
| | *bglu3-2* | RS1847<br>T35S | TTATTCACCAAGCAAACTCTTGAA<br>TCTGGGAACTACTCACAC |
| | *bglu4-2* | RS1259<br>3144 | ATGGAACAGATTTTGGCTCTGTTTGCCATTTT<br>GTGGATTGATGTGATATCTCC |
| | *bglu4-2* | 3144<br>JFF001 | GTGGATTGATGTGATATCTCC<br>CATCATTCTGTGGTTGAGCCATCC |
| cDNA cloning | *BGLU1* | RS1385<br><br>RS1386 | GGGGACAAGTTTGTACAAAAAAGCAGGCTCCA<br>TGGA AGATGTTTTGACTCTC<br>GGGGACAAGTTTGTACAAGAAAGCTGGGTATC<br>TGGA AGAAGAGAAGTTGCTATGC |
| | *BGLU3* | RS1387<br><br>RS1388 | GGGGACAAGTTTGTACAAAAAAGCAGGCTCCA<br>TGGAACTGACTTTGTCTCTG<br>GGGGACCACTTTGTACAAGAAAGCTGGGTAC<br>GAAGAAGCTGAAGAAGAAAAG |
| | *BGLU4* | RS1255<br><br>RS1256 | GGGGACAAGTTTGTACAAAAAAGCAGGCTCAA<br>TGGA ACAGATTTTGGCTCTGTTTGCC<br>GGGGACCACTTTGTACAAGAAAGCTGGGTCG<br>AAGGA AGGAGAGAGGTTGCTCTGC |

| | | | |
|---|---|---|---|
| qRT-PCR | *BGLU1* | JFF032<br>JFF033 | GCCGGGATATCTGCTTATCA<br>CAAGCTATGTCTCCATTATCCATTT |
| | *BGLU3* | JFF059<br>RS1320 | GGTCTAGGCTTATACCGAATGG<br>AGTTCCCTGATGAGCAGTTTCTACC |
| | *BGLU4* | JFF072<br>JFF073 | ACTACCCTGAGGGATTCG<br>GTGGCTTACTAACTCTTGG |
| | *PEROXIN4* | RS935<br>RS936 | TTGGACGCTTCAGTCTGTGT<br>TGAACCCTCTCACATCACCA |
| | *EF1α* | JFF028<br>JFF029 | ACCACCACTGGGCACTTGATC<br>GTGCAGTAGTACTTGGTGGTCT |
| RT-PCR | *BGLU1* | RS1302<br>RS1390 | ACAGCAGGAGCGATTTCCCGGAAG<br>CCAAATCTTGGTTCATTGTTTTACC |
| | *BGLU3* | RS1319<br>JFF010 | GAGATGCAGCGACAAGAACGACTTCC<br>ACGAAGAAGCTGAAGAAGAAAAGTTGCTCTGC |
| | *BGLU4* | RS1260<br>RS1261 | CTAGAAGGAAGAAGAGTATTTGCTCTGCA<br>CAACCCAAAGAGCCCAAGATTTCTA |
| | *ACTIN2* | RS469<br>RS470 | TCCGCTCTTTCTTTCCAAGCT<br>TCCAGCACAATACCGGTTGTA |
| Overexpression genotyping | *2x35S::BGLU1* | RS1434<br>RS1390 | GAATTCATCGCCCGGGAC<br>CCAAATCTTGGTTCATTGTTTTACC |
| | *2x35S::BGLU3* | RS1434<br>RS1320 | GAATTCATCGCCCGGGAC<br>AGTTCCCTGATGAGCAGTTTCTACC |
| | *2x35S::BGLU4* | RS1434<br>RS1473 | GAATTCATCGCCCGGGAC<br>AAGCCGCATGATAGTGTATGAC |
| Transgene overexpression | *2x35S::BGLU1* | RS1516<br>RS1320 | CCGTTCCATGGGCTAGAAGCTTCTCCTCC<br>AGTTCCCTGATGAGCAGTTTCTACC |
| | *2x35S::BGLU3* | RS1516<br>RS1474 | CCGTTCCATGGGCTAGAAGCTTCTCCTCC<br>TGGCAGTCTTGATCCAATGGTTCTTTTC |
| | *2x35S::BGLU4* | RS1516<br>RS1474 | CCGTTCCATGGGCTAGAAGCTTCTCCTCC<br>TGGCAGTCTTGATCCAATGGTTCTTTTC |

**Table S2. Oxford Nanopore Technologies sequencing data sets.** *bglu* mutant IDs, ENA IDs, the numbers of reads and nucleotides from the long-read sequencing run, the chosen flow cells and the sequences across both T-DNA::genome junctions are given. The sequences are shown with the genomic sequences given in bold, the T-DNA sequences given in normal letters and sequences that go back to the insertion event and neither match genomic *BGLU* or T-DNA sequences given in italics.

| T-DNA insertion mutant ID (gene ID) | ENA ID | Read number | Nucleotides | Flow cell | T-DNA::genome junction sequences |
|---|---|---|---|---|---|
| ***bglu1-1*** GABI_341B12 (*At1g45191*) | ERS4255859 | 1,014,594 | 13,523,523,050 | R9.4.1 | RB at position 17,116,579<br>**CACGTAATG***AAGACAATATGGGAGACTAGTCTCCCT CATACTCATTGTATGTCCATATGGGAGACTAGTC*TC CAGTCGG<br>LB at position 17,116,598<br>CCATTTACA*TGAGTACCA***TATTTTGTT** |
| ***bglu3-2*** GABI_853H01 (*At4g22100*) | ERS4255860 | 804,401 | 2,736,092,996 | R9.4.1 | LB at position 11,709,058<br>**ATGGAGGAT***AAATCATC*TATATTCAA<br>RB at position 11,708,997<br>AGTGTTTGA*TATAATTCTTC***TCATGCTTC** |
| ***bglu4-2*** SALK_029729 (*At1g60090*) | ERS4255861 | 1,308,191 | 9,914,725,163 | R9.4.1<br>R10 | LB at position 22,156,000<br>**GCTTCTCTG**AACAAATTG<br>LB at position 22,156,017<br>TCAATTTGT*CCGCAATGTGTTGTTGTCT***ATGAGGTTT** |

**Table S3. BGLU3 candidate features in ESI$^+$ mode.** The metabolic features with their mass-to-charge ratios (*m*/*z*) and retention times (RT) after alignment across samples are listed, followed by putative ion types with charge states as well as the *m*/*z* values of the MS/MS fragments (at the indicated collision energies). The *m*/*z* of the precursors may slightly differ from the *m*/*z* of the feature names, as MS/MS spectra were taken from single samples. Only precursors and fragments with an intensity of at least 50 counts and 5% relative intensity are listed; if there were more than 15 of them, only those 15 ones with the highest intensities are shown. Putative isotopes were removed from the list; however, *m*/*z* pointing to isotopes but with an unexpected high intensity that could indicate an overlap with another fragment were included. For doubly charged fragments, the isotope series are given in parentheses. Precursors are given in brackets, dominant fragments (≥ 30% intensity) are highlighted in bold. Fragments, which we refer to in the Results and discussion part and that occurred in the MS/MS spectra but at < 5% intensity, as well as isotopes of doubly charged fragments with < 5% intensity but > 50 counts are given in gray and in italics. For the doubly charged precursors, peaks with *m*/*z* values higher than those of the precursors were also considered (see underlined *m*/*z*), as they may represent fragments with single charges. Moreover, for the doubly charged precursors, the charges of the fragments are given, if they could be derived from the isotope pattern. For doubly charged fragments one decimal place is given, while for the other fragments nominal masses are given.

<table>
<tr><th>Candidate products and substrates</th><th>Feature m/z</th><th>Feature RT (min)</th><th>Putative ion type with charge state</th><th>Collision energy and MS/MS fragments (m/z)</th></tr>
<tr><td>Main products</td><td>866.4164</td><td>11.75</td><td>[M]$^+$</td><td>**37.3 eV:** **822**, 640, *398*, *339*, 289, 207, **104**</td></tr>
<tr><td></td><td></td><td></td><td></td><td>**60.0 eV:** 822, 640, 520, 399, **398**, 374, **339**, 338, 337, 332, 267, 253, 251, **104**, 99</td></tr>
<tr><td></td><td>433.7128</td><td>11.76</td><td>[M+H]$^{2+}$</td><td>**28.3 eV:** *398*$^{1+}$, (*271.7* | *271.2* | *270.7* | *270.2* | *269.7* | **269.2**)$^{2+}$, (*261.7* | *261.2* | *260.7* | 260.2)$^{2+}$, (*201.1* | *200.6* | *200.1* | *199.6* | *199.1* | **198.6**)$^{2+}$, 140$^{1+}$, 85$^{1+}$</td></tr>
<tr><td></td><td>773.2116</td><td>10.51</td><td>[M+H]$^+$</td><td>**35.0 eV:** 449, **303**, 145, 127, 85</td></tr>
<tr><td>Main substrates</td><td>704.3635</td><td>13.11</td><td>[M]$^+$</td><td>**34.1 eV:** **660**, 478, 398, *339*, **104**</td></tr>
<tr><td></td><td>330.6915</td><td>13.11</td><td>[M+H–$CO_2$]$^{2+}$</td><td>**25.8 eV:** (*269.7* | 269.2 | *268.7* | *268.1*)$^{2+}$, 251$^{1+}$, (*199.6* | *199.1* | **198.6** | 198.1 | 197.6)$^{2+}$, (*170.6* | *170.1* | *169.6* | 169.1 | *168.1*)$^{2+}$, 140, 122</td></tr>
</table>

| | | | | |
|---|---|---|---|---|
| | 781.3270 | 15.38 | $[M+H]^+$ or $[M]^+$ | **35.6 eV:** 619, **601**, 575, 496, 480, 478, 460, **452**, 339, 337, 311, 289, 287, 271, 166 |
| Minor products | 411.7178 | 11.87 | $[M+H]^{2+}$ | **30.0 eV:** 398$^{1+}$, (269.7 \| 269.2)$^{2+}$, (*261.2* \| *260.7* \| 260.2)$^{2+}$, 207$^{1+}$, (*200.1* \| *199.6* \| *199.1* \| **198.6**)$^{2+}$, 175$^{1+}$, 169, 140, 97, 85 |
| | 432.7049 | 14.62 | $[M+H]^{2+}$ | **30.0 eV:** 372$^{1+}$, (*330.7* \| *330.2* \| 329.7)$^{2+}$, 287$^{1+}$, (*261.1* \| *260.7* \| *260.1* \| *259.6* \| **259.1**)$^{2+}$, (*239.6* \| 239.1 \| *238.6* \| 238.1)$^{2+}$, (*230.6* \| *230.1* \| 229.6)$^{2+}$, 227, 148$^{1+}$, 140$^{1+}$, 127$^{1+}$, 122$^{1+}$, 97, 85$^{1+}$ |
| | 440.7207 | 12.45 | $[M+H]^{2+}$ | **30.0 eV:** 424$^{1+}$, (*298.6* \| 298.2)$^{2+}$, (*282.6* \| 282.1)$^{2+}$, (*274.1* \| *273.6* \| **273.1**)$^{2+}$, 252, (*229.1* \| **228.6** \| *228.1* \| **227.6** \| *227.1*)$^{2+}$, (*213.1* \| **212.6** \| *212.1* \| **211.6**)$^{2+}$, **140**, 127,122, 97, 85 |
| | 440.7209 | 12.90 | $[M+H]^{2+}$ | **28.5 eV:** 424$^{1+}$, (*344.2* \| 343.7)$^{2+}$, (*283.7* \| *283.2* \| 282.6 \| **282.1**)$^{2+}$, (*274.7* \| *274.1* \| 273.6 \| **273.1**)$^{2+}$, (*229.1* \| 228.6 \| *228.1* \| **227.6** \| *227.1*)$^{2+}$, (*213.6* \| *213.1* \| **212.6** \| *212.1* \| **211.6**)$^{2+}$, 176$^{1+}$, **140**$^{1+}$, 127, **122**, 85 |
| | 441.7099 | 9.45 | $[M+H]^{2+}$ | **30.0 eV:** 553$^{1+}$, 413$^{1+}$, (*278.2* \| *277.7* \| 277.2)$^{2+}$, 267$^{1+}$, (259.6 \| 259.1)$^{2+}$, 225$^{1+}$, (*207.6* \| *207.1* \| **206.6**)$^{2+}$, (*193.1* \| 192.6)$^{2+}$, 145, 142, 140$^{1+}$, 127, **124**$^{1+}$, 85 |
| | 441.7102 | 9.96 | $[M+H]^{2+}$ | **30.0 eV:** 553$^{1+}$, 390$^{1+}$, (*330.2* \| 329.7)$^{2+}$, 287$^{1+}$, (*277.7* \| 277.2)$^{2+}$, **267**$^{1+}$, (*260.1* \| *259.6* \| **259.1**)$^{2+}$, (208.1 \| *207.6* \| *207.1* \| **206.6**)$^{2+}$, 148, 140$^{1+}$, **124**$^{1+}$, 122, 97, **85** |
| | 448.7181 | 11.05 | $[M+H]^{2+}$ | **30.0 eV:** (*366.2* \| *365.7* \| 365.2)$^{2+}$, 287$^{1+}$, (*285.2* \| *284.7* \| 284.2)$^{2+}$, (*260.1* \| *259.6* \| 259.1)$^{2+}$, (*215.1* \| *214.6* \| *214.1* \| **213.6**)$^{2+}$, (*207.1* \| 206.6)$^{2+}$, 140$^{1+}$, 127, 124, 85 |
| | 460.1686 | 11.76 | $[?]^{2+}$ | **30.0 eV:** 397$^{1+}$, 396$^{1+}$, 357, 339, (*305.6* \| **305.1**)$^{2+}$, (*297.6* \| *297.1* \| *296.6* \| 296.1 \| 295.6 \| *295.1*)$^{2+}$, (*288.1* \| *287.6* \| **287.1** \| 286.6 \| *286.1* \| *285.6* \| *285.1*)$^{2+}$, (*282.1* \| *281.6* \| **281.1** \| 280.6)$^{2+}$, 258, 237, 104 |

| | | | | |
|---|---|---|---|---|
| | 467.1764 | 12.89 | $[M+H]^{2+}$ | **30.0 eV:** (*371.1* \| **370.6**)$^{2+}$, (334.1)$^{2+}$, (*325.6* \| **325.1**)$^{2+}$, **318**, (316.6 \| 316.1)$^{2+}$, (*310.6* \| **310.1** \| **309.6** \| **309.1**)$^{2+}$, (*301.1* \| **300.6** \| **300.1**)$^{2+}$, 160, **142**, 130, 85 |
| | 494.1968 | 19.11 | $[M+H]^{2+}$ | **30.0 eV:** 452$^{1+}$, 339$^{1+}$, 337$^{1+}$, 311$^{1+}$, **207**$^{1+}$, **175**$^{1+}$, 147 |
| | 536.7408 | 17.07 | $[M+H]^{2+}$ | **30.0 eV:** 312, 294$^{1+}$, 245, **207**$^{1+}$, 175$^{1+}$ |
| | 757.2152 | 11.63 | $[M+H]^{+}$ or $[M]^{+}$ | **35.0 eV:** 433, **287**, 145, 85 |
| | 757.2166 | 11.42 | $[M+H]^{+}$ or $[M]^{+}$ | **35.0 eV:** 449, **303**, 147, 129, 85 |
| | 943.3804 | 13.74 | $[M+H]^{+}$ or $[M]^{+}$ | **40.0 eV:** 781, **763**, **737**, 658, 640, **614**, 601, 575, 475, 452, **449**, 337, 331, 289, **287** |
| Minor substrates | 287.1382 | 7.25 | $[M+H]^{+}$ or $[M]^{+}$ | **25.0 eV:** [287], **270**, 269, **242**, **148**, **147**, **146**, **140**, 122, **119**, **118** |
| | 402.1451 | 13.11 | $[?]^{2+}$ | **30.0 eV:** 309, 305, (*297.1* \| **296.6** \| **296.1**)$^{2+}$, 294, (*288.1* \| **287.6** \| **287.1** \| **286.6**)$^{2+}$, (*281.6* \| **281.1**)$^{2+}$, 280, (**223.6** \| **223.1**)$^{2+}$, (**214.6** \| *214.1*)$^{2+}$, **130**, **104** |
| | 575.2851 | 18.19 | $[M+H]^{+}$ or $[M]^{+}$ | **30.0 eV:** **452**, 313, 311, 287, 148 |
| | 619.2742 | 17.37 | $[M+H]^{+}$ or $[M]^{+}$ | **32.4 eV:** 601, 480, 478, 460, **452**, 357, **339**, 337, 313, 311, 289, 287, 271, 166, 140 |
| | 633.2899 | 19.68 | $[M+H]^{+}$ or $[M]^{+}$ | **35.0 eV:** **601**, 510, **478**, **460**, **371**, 369, **339**, **338**, **337**, 313, 297, 287, 271, 166, 140 |
| | 660.3742 | 13.21 | $[M]^{+}$ | **35.0 eV:** [**660**], 538, 520, 515, 478, 398, 374 |
| | 795.3419 | 17.03 | $[M+H]^{+}$ or $[M]^{+}$ | **37.0 eV:** 633, **601**, 510, 478, 460, 369, 337, 287, **207**, 206, 175 |

**Table S4. Putative modifications explaining further BGLU3 candidate products in ESI$^+$ mode.** The names of the metabolic features including mass-to-charge ratios (*m*/*z*) and retention times (RT) derived from alignment across samples are shown. Only features are listed that may represent metabolites derived from the metabolite belonging to the feature with an *m*/*z* value of 866.4164 at 11.75 min. For calculations of the neutral masses of the metabolites, it was assumed that the feature with an *m*/*z* value of 866 (given in bold) is an $[M]^+$ ion and that the other features are doubly charged, representing $[M+H]^{2+}$ ions (see Supplementary Table S3). Differences of the neutral masses to that of the feature with an *m*/*z* value of 866 are given and putative modifications fitting to these differences are indicated. The features are sorted according to increasing *m*/*z*.

| Feature (*m*/*z*, RT) | Neutral mass (Da) | Mass difference (Da) | Putative modification |
|---|---|---|---|
| ***m*/*z* 866.4164, 11.75 min** | **865.4** | | |
| *m*/*z* 411.7178, 11.87 min | 821.4 | -44 | $-CO_2$ (decarboxylation) |
| *m*/*z* 432.7049, 14.62 min | 863.4 | -2 | -2H (dehydrogenation) |
| *m*/*z* 440.7207, 12.45 min | 879.4 | +14 | $+CH_2$ (methylation) |
| *m*/*z* 440.7209, 12.90 min | 879.4 | +14 | $+CH_2$ |
| *m*/*z* 441.7099, 9.45 min | 881.4 | +16 | +O (hydroxylation) |
| *m*/*z* 441.7102, 9.96 min | 881.4 | +16 | +O |
| *m*/*z* 448.7181, 11.05 min | 895.4 | +16 +14 | +O $+CH_2$ |
| *m*/*z* 467.1764, 12.89 min | 932.3 | +52.9 +14 | + unknown group 1 (UG1) $+CH_2$ |
| *m*/*z* 494.1968, 19.11 min | 986.4 | +52.9 +14 +16 + (2*18) +2 | +UG1 $+CH_2$ +O $+2H_2O$ +2H |
| *m*/*z* 536.7408, 17.07 min | 1071.5 | +206 | unknown |

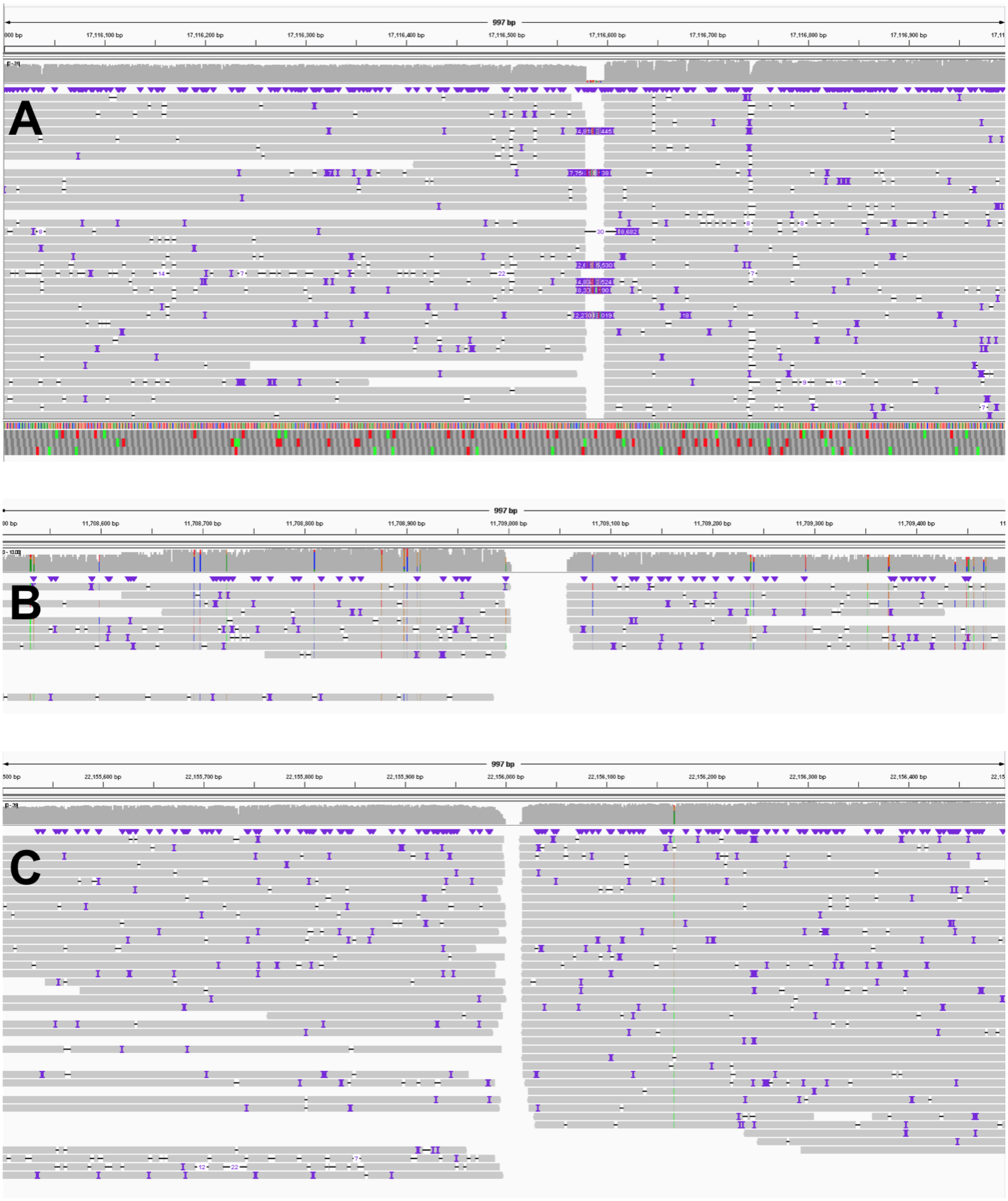


**Fig. S1.** Oxford Nanopore Technologies long-read sequencing of T-DNA insertion mutants. The screenshots of the long-read sequencing results, mapped to the Col-0 genome sequence, show homozygosity of the T-DNA insertion mutants. (A) *bglu1-1*, (B) *bglu3-2* and (C) *bglu4-2*, gaps indicate where the T-DNA insertions are located.

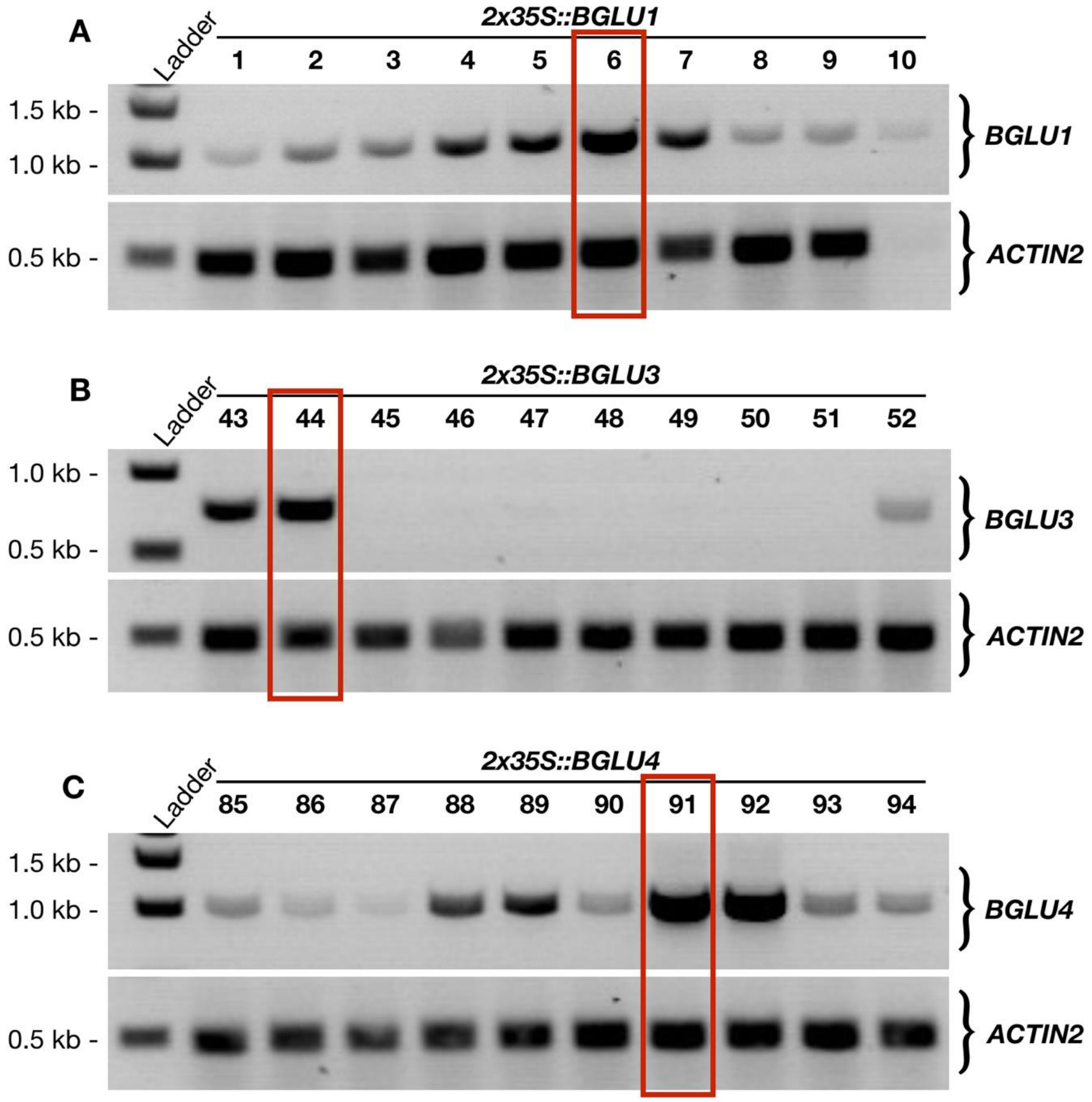


**Fig. S2.** Transgene expression in *2x35S::BGLU* lines. Transgene expression was examined by RT-PCR in rosette leaves of *A. thaliana* and is shown for ten lines of (A) *BGLU1*, (B) *BGLU3* and (C) *BGLU4*. The red rectangles indicate the lines that were used for further experiments.

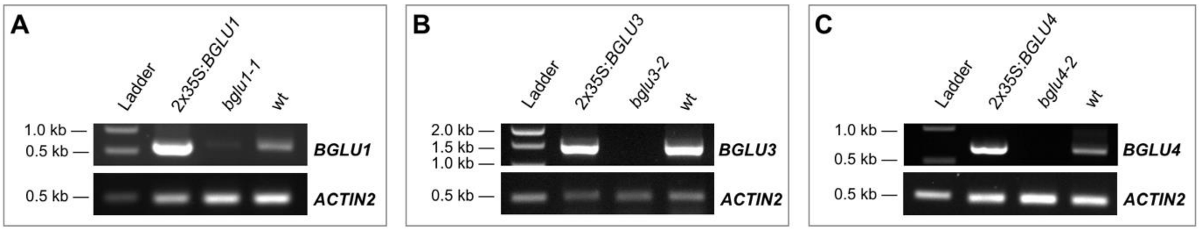


**Fig. S3.** Transcript levels of *BGLU1*, *BGLU3* and *BGLU4* in the expression variant lines. RT-PCR results for the intact transcripts of *A. thaliana* (A) *BGLU1* in rosette leaves, (B) *BGLU3* in dry mature seeds and (C) *BGLU4* in mature seeds, which were soaked 24 h in water, are shown. For each *BGLU* gene, the *2x35S::BGLU* line, the *bglu* mutant and the Col-0 wild type (wt) were analyzed. *ACTIN2* was used as reference.
